# Predicting human prediction error empowers reward learning task design

**DOI:** 10.1101/2024.12.08.627441

**Authors:** Jaehoon Shin, Jee Hang Lee, Sang Wan Lee

**Affiliations:** Department of Bio and Brain Engineering, Korea Advanced Institute of Science and Technology (KAIST) 34141 Daejeon, Republic of Korea; Department of Human-Centered Artificial Intelligence, Sangmyung University 03016 Seoul, Republic of Korea; Institute for Advanced Intelligence Study, Daejeon, Republic of Korea; KAIST Center for Neuroscience-inspired Artificial, KAIST, Daejeon, 34141, Republic of Korea; Department of Brain and Cognitive Sciences, Korea Advanced Institute of Science and Technology (KAIST); Graduate School of Data Science, Korea Advanced Institute of Science and Technology (KAIST); Kim Jaechul Graduate School of AI, Korea Advanced Institute of Science and Technology (KAIST)

## Abstract

Environmental conditions affect human reward prediction. Stable environments foster accurate prediction but constrain learning opportunities, whereas uncertain environments diminish predictability. This stability-uncertainty dilemma complicates task design. We conceptualize this challenge as a task learning paradigm termed meta-prediction – predicting human prediction itself. The meta-prediction entwines two Bellman equations: one emulating human reward learning while the other generates new tasks by predicting the prediction error arising from the first. The meta-prediction with 82 subjects’ data generated subject-independent tasks across four distinct scenarios. These tasks orchestrate foraging and uncertainty conditions, confirming our framework’s task design ability. Moreover, their mechanistic interpretability provides insight into human reward learning. An independent fMRI study with 49 individuals validated that these tasks effectively modulated behavior and neural activities in prediction error encoding regions, including ventral striatum and lateral prefrontal cortex. Lastly, we demonstrated its compositional capacity to generate complex tasks, uncovering intrinsic biases in human reward learning.

## Introduction

The environmental conditions exert a profound influence on human reward learning. Previous research has yielded valuable insights into reward learning, including learning action values for habit formation^1–4^ and using internal models for goal-directed learning^5–8^. These investigations have primarily relied on a stable environmental setting, which enables confident predictions. While these investigations have predominantly occurred within stable environmental contexts that foster accurate predictions, they inadvertently constrain the breadth of experience and limit the predictive capabilities.

Recent studies have leveraged volatile and uncertain environments to examine the nuanced dynamics of reward prediction^−^^12^. Such approaches accommodate the view that learning in complex environments demands sophisticated predictive strategies wherein complementary functions— habit, goal, memory, and intuition^13,14^— work in concert. However, the efficacy of such strategies can be compromised in a highly uncertain and volatile environment, making reliable predictions difficult. This tension reveals a key dilemma in task design: both environmental stability and uncertainty impose constraints on our reward learning.

To address this, we conceptualize this challenge as a task learning problem (decoding), which incorporates how environmental stability and uncertainty shape human reward prediction processes (encoding). This perspective leads to a novel solution: learning a task control policy that predicts human prediction, a paradigm we term *meta-prediction* (Fig. 1A). The meta-prediction applies the reward prediction principle in machine learning, formulated as Bellman updates^15^, to human reward prediction, which can likewise be specified as Bellman updates (Fig. 1B). Unlike conventional task design and optimization schemes^16,17^, this setting enables us to generate diverse learning tasks that guide the human reward prediction in targeted ways (Fig. 1C: encoding-decoding framework).

**Fig 1.**
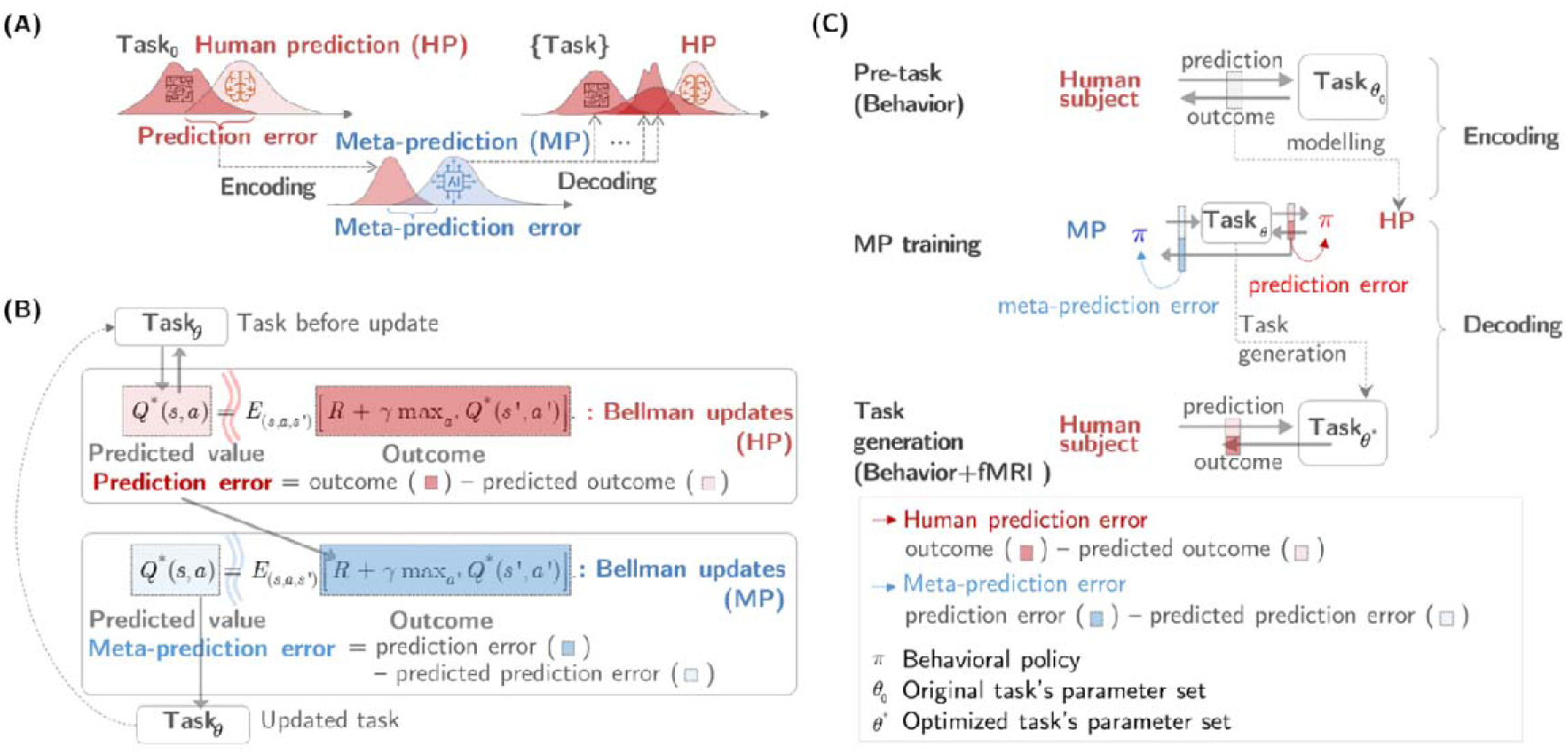
Meta-prediction. **(A)** Meta-prediction problem. (Left) In an uncertain value-learning environment, the human prediction model (HP) aims to learn the probability distribution of the task’s reward and structure by minimizing the mismatch between the estimated and true distribution, called prediction error. (Middle) Meta-prediction model (MP) aims to learn the distribution of individual HP errors from the HP-task interaction (*encoding)*. (Right) Once learning is completed, meta-prediction can generate the task by changing the task parameters to guide HP (*decoding*). **(B)** Formulating the MP problem as a coupled Bellman equation. The overall framework consists of the two-stage predictions, each specified as a separate Bellman equation. The red and blue correspond to HP and MP, respectively. MP’s reward function is defined as the prediction error of the HP. The MP’s state space is defined as the task variable and the HP’s prediction error. The MP’s actions are defined as a set of changes of the task parameters. The MP’s prediction error (i.e., “meta-prediction error”) is defined as the difference between the expected and the actual prediction error of the HP. The MP problem can be solved using reinforcement learning (RL) algorithms. **(C)** The meta-prediction framework comprises three steps: pre-task for data acquisition, HP fitting, and MP training. First, human choice behavior data were collected using a reward learning task, such as a two-step goal-directed learning task^8,10^. Second, HPs were fitted to imitate the individual subject’s behavior (“encoding”). Third, another RL model (MP) was trained to change the task parameters so as to guide the key variables of the individual HP. This training process yields an MP that generates a task design to guide a single subject’s behavior (“decoding”). Note that our framework is applicable to any parameterized task.

Leveraging the fact that the prediction error—defined as the discrepancy between human predictions and actual environmental outcomes—serves as a readout of the interaction between the environmental structure and human predictions, we trained the meta-prediction framework to generate tasks that either minimize or maximize human prediction error. Extremizing this signal creates a dynamic space in which learners navigate the tradeoff between environmental stability and uncertainty. For precise modulation of reward learning strategies, the framework targets the two distinct forms of prediction error known to guide habitual and goal-directed learning^2,6,10,18^: reward prediction error (RPE) and state prediction error (SPE).

To demonstrate the feasibility of the meta-prediction framework, first, we obtained subject-specific human prediction models by fitting the computational model of RL to behavioral data from 82 subjects in a 2-stage goal-directed learning task^7,8,10^. Second, we trained the meta-prediction module (“meta-prediction” of Fig. 1) to minimize or maximize SPE and/or RPE of the pre-trained human prediction model. These generated tasks were verified in two independent fMRI studies with a total of 49 participants. The behavior analyses examined whether and how these interventions influence subjects’ performance and learning speed. The model-based fMRI analyses demonstrated the intervention effects on the neural activities of brain areas known to guide predictions

## Result

### Meta-prediction theory

Our core challenge is to design tasks accommodating the ways in which environmental stability and uncertainty shape human reward prediction. The proposed framework, termed *meta-prediction*, addresses this problem by systematically generating tasks based on predictions of human prediction.

The meta-prediction consists of two steps. It learns to predict human predictions about future outcomes (*human prediction* in Fig. 1A) and then uses this prediction to generate a new task design (meta-prediction in Fig. 1A). Importantly, this design ensures that meta-prediction generates a task design that effectively influences *human prediction*, thereby enhancing environmental diversity without losing learnability. Our framework is generalizable in that one can generate various task designs by applying different task optimization schemes. For example, one can make a curriculum easier by having the meta-prediction minimize the human prediction error, or alternatively, make the existing task more challenging by maximizing it. Moreover, the meta-prediction allows us to diversify the dimensions of scenarios by incorporating into meta-prediction various factors, such as individual differences (e.g., varying the parameters of the human prediction model), type (e.g., state or reward), or the profile of prediction (e.g., reducing, maintaining, or increasing the amount of prediction error).

To illustrate the theoretical background, our framework was designed based on the Bellman equation, the basic mathematical framework of reward prediction. Weaved into a single learning framework are two Bellman equations, one for human prediction and the other for meta-prediction (Fig. 1B and Supplementary Table 4-6 for algorithm description). The Bellman equation for human prediction learns to predict rewards of the environment (task) while the Bellman equation for meta-prediction learns to predict the residual of human prediction (prediction error) by changing the task variables. The role of the former is to provide full observations about latent variable changes of the human prediction process, so that the latter can effectively translate them into the task design. This completes a learning cycle of “task - human prediction - meta-prediction - task updates”.

The learning cycle begins with individually fitted human prediction models to encode individual variability of human prediction strategies (the *encoding* step in Fig. 1C). To preclude overfitting issues, these models should adopt theoretical principles from neuroscientific studies^6–8^ rather than data-driven approaches using a generic class of neural networks. The model used in this study accommodates the prefrontal control of model-based and model-free reinforcement learning^8^. Then, we train another learning agent, called a meta-prediction model, to optimize the task design by predicting human prediction error, estimated from the human prediction module (*decoding* step in Fig. 1C). The meta-prediction network was implemented with a deep reinforcement learning model^19,20^.

### Fully parameterized task space with varying predictability

To furnish the meta-prediction framework with a broad and flexible task space, we generalized a conventional two-step Markov decision task^7,8,10^. The task paradigm was carefully parameterized to accommodate stable goal-directed learning and adaptive foraging with different degrees of uncertainty (Fig. 2A). Unlike traditional reward learning regimes in which reward functions remain fixed regardless of subjects’ choices to impose rigid constraints on the human learner’s prediction, our task parameters were crafted to support a wide spectrum of predictability (Fig. 2B). They include (1) state-transition probability, which directly influences the learner’s prediction about state-transition, (2) goals, which links the learner’s prediction to reward, and (3) reward foraging, in which the reward changes as a function of the learner’s action.

**Figure 2.**
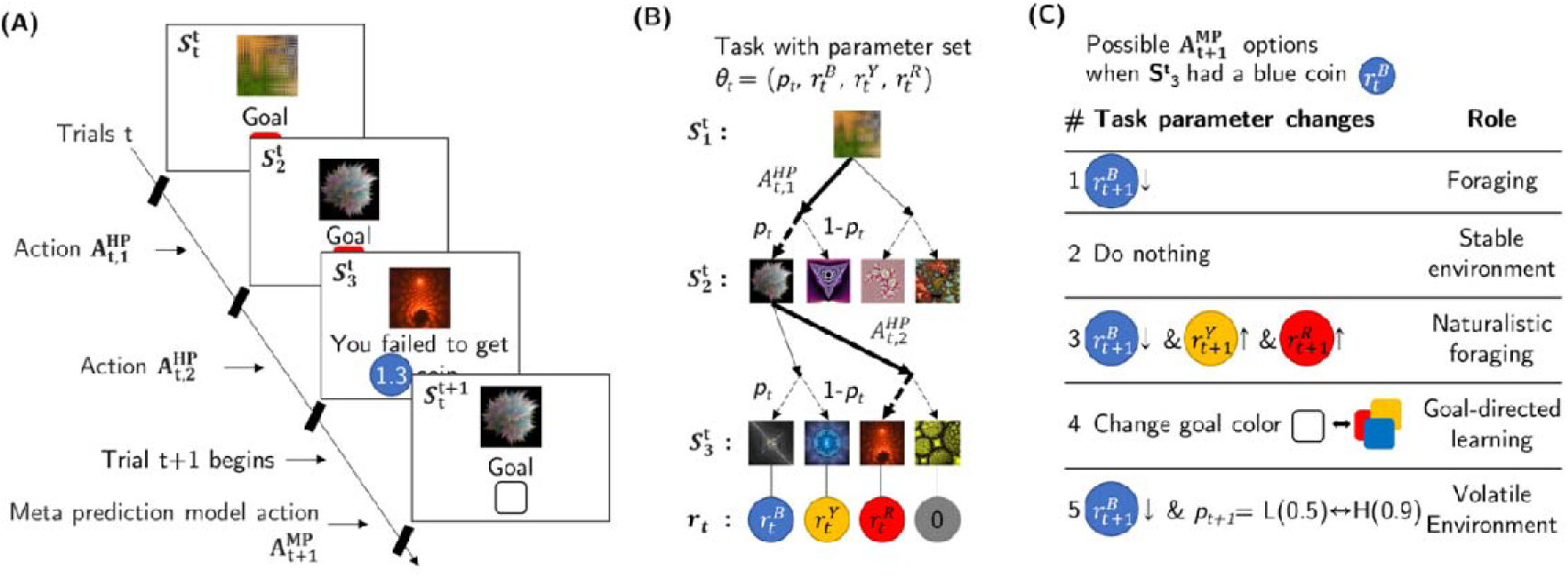
Parameterized reward learning task. **(A)** Markov decision task^8,10^ for goal-directed learning and foraging. This task accommodates goal-directed learning and foraging under environmental uncertainty. The learner, either a human subject or a human prediction model (HP), makes two consecutive choices in a single trial to collect coins of different colors. In each trial, subjects are presented with one of the three goals indicated by different colors. Subjects are asked to collect a coin available at the outcome state. Each outcome state contains a coin of a specific color, associated with a different reward amount. Coins are redeemable only when the current goal color matches the coin color, enforcing goal-directed learning. If the goal color is blank, all coins are redeemable, making the task equivalent to conventional reward learning. To reach the desired outcome state, subjects must learn to navigate the state-space whose structure is determined by state-transition probabilities. The fractal image and a box represent a state and a goal, respectively. **(B)** The structure of the task. The task parameters (θ_t_) consist of the state transition probability (*p_t_*) and the reward associated with different coin colors (r_t_). The reward amount r_t_ may change over time by the associated Meta-prediction (MP) model’s actions. **(C)** The MP’s actions are defined on the task parameter space. Each action has a distinctive role: foraging (action #1, decreasing the reward in the visited goal state), stable environment (action #2, preserving the current task environment setting), naturalistic foraging (action #3, decreasing the visited goal state’s reward while recovering other goal states’ rewards), goal-directed learning (action #4, changing the target goal states), volatile environment (action #5, switching the state-transition probability between 0.9 and 0.5).

First, for the state-transition, we used a switching probability^8^ instead of a drifting probability^10^ because the former exerts a more dramatic impact on state prediction. The transition probability switches between 0.9 and 0.5, each simulating the condition of low and high predictability about state-transition, respectively. Second, the goal conditions were adapted from Lee et al.(2014)^8^, in which the reward is redeemable only if the goal and the color of the coin obtained at the outcome state match (goal-dependent devaluation). The goal is assigned at the beginning of each trial to facilitate reward prediction and goal-directed behavior. Lastly, we introduced reward foraging (Fig. 2C). It is our major modification to the existing two-step task paradigm. Here, the learner’s reward acquisition decreases reward by a certain ratio while rewards in the unvisited outcome states are gradually restored, accommodating foraging in nature. Note that this action-dependent reward function creates complex dynamics of the learner’s predictions about future reward.

Our meta-prediction model adjusted these task parameters, each designed to serve a distinct purpose: (1) foraging environment (decreasing the reward in the exploited goal state), (2) stable environment (preserving the current task structure, maintaining environmental consistency), (3) naturalistic foraging environment (reducing the exploited goal state’s reward while replenishing rewards in previously unexploited goal states), (4) goal-directed learning environment (explicitly assigning target goal states), (5) volatile environment (altering the state-transition probability between 0.9 and 0.5, introducing different level of uncertainty) (Also refer to the table of Fig. 2C). Importantly, each task parameter adjustment corresponds to a discrete action of the meta-prediction model’s control policy. For example, the model’s action sequence 2-1-3-3 represents a specific task design: the environment begins in a stable state (Action 2), transitions into reward depletion through foraging (Action 1), and evolves into a more naturalistic foraging pattern (Action 3s), where rewards shift adaptively between exploited and unexploited goal states.

Our parameterized task space enables the meta-prediction model to strategically intervene in human prediction processes by shaping the structure and dynamics of the learning environment. For example, if the meta-prediction model predicted the moment the learner predicts reward acquisition, it would determine when and how to intervene in its prediction by changing the reward function. Likewise, the meta-prediction model predicting a human’s prediction about state-transition could engage in human prediction by changing the structure of the state space.

### Meta-prediction networks

We trained the *meta-prediction* framework on the task parameter space defined in the preceding section. The training process comprised two stages (illustrated in Fig. 1A): In the encoding stage, the “*human prediction”* model (termed as HP) was trained to predict the human’s choice behavior. In the decoding stage, the “*meta-prediction”* models (termed as MP) were trained to generate new tasks, using the *human prediction* model as a reward function.

The HP was implemented using a reinforcement learning algorithm that combines model-based and model-free reinforcement learning on a trial-by-trial basis, which is known to explain goal-directed and habitual reward learning^8^. Importantly, the HP affords the MP the leverage to predict human prediction error, the key variable guiding human prediction. The HPs’ parameters were optimized to maximally match their value function with the choice behavior of 82 individual human subjects, collected using the two-step Markov decision task^8,21^. Model comparison and parameter recoverability analysis in the prior studies^8,22,23^ ensured the model’s reliability in predicting human reward learning.

We then trained the MP (the learning agent) to learn the HP’s prediction (the reward function). To make the MP fully compatible with the HP, the state space of the MP included the HP’s state and/or reward prediction error, which is the key variable for model-based and model-free learning (see the ‘Meta-prediction models’ section in the Method). The MP’s action refers to changing the task parameters (Fig. 2B). Accordingly, a single learning cycle of the MP is illustrated as follows. The MP’s action is followed by the HP’s prediction, which is followed by the MP’s observation about the HP’s prediction error. The MP then computes the discrepancy between predicted and actual prediction errors of the HP (*meta-prediction error* in Fig. 1B), based on which the MP updates its value function. The MP was implemented using a double deep Q network.^19,20^

The MP was trained in four training conditions: maximizing/minimizing x reward/state prediction error of HPs (MaxR, MinR, MaxS, and MinS; see Fig. 3A). In MaxR/MinR, the MP learns to predict the HP’s current absolute reward prediction errors, whereas in MaxS/MinS, the MP aims to predict the HP’s state prediction errors. For MaxR/MaxS, the reward function for the MP was defined as being directly proportional to the HP’s prediction error. Conversely, in MinR/MinS, the reward function was inversely proportional to the HP’s prediction error. Note that these training settings achieve the key challenge of our *meta-prediction* problem: by manipulating environment stability and uncertainty (corresponding to MP’s policy), the MP can effectively ensure a wide spectrum of predictability of the HP (quantified as MP’s reward functions).

**Fig 3.**
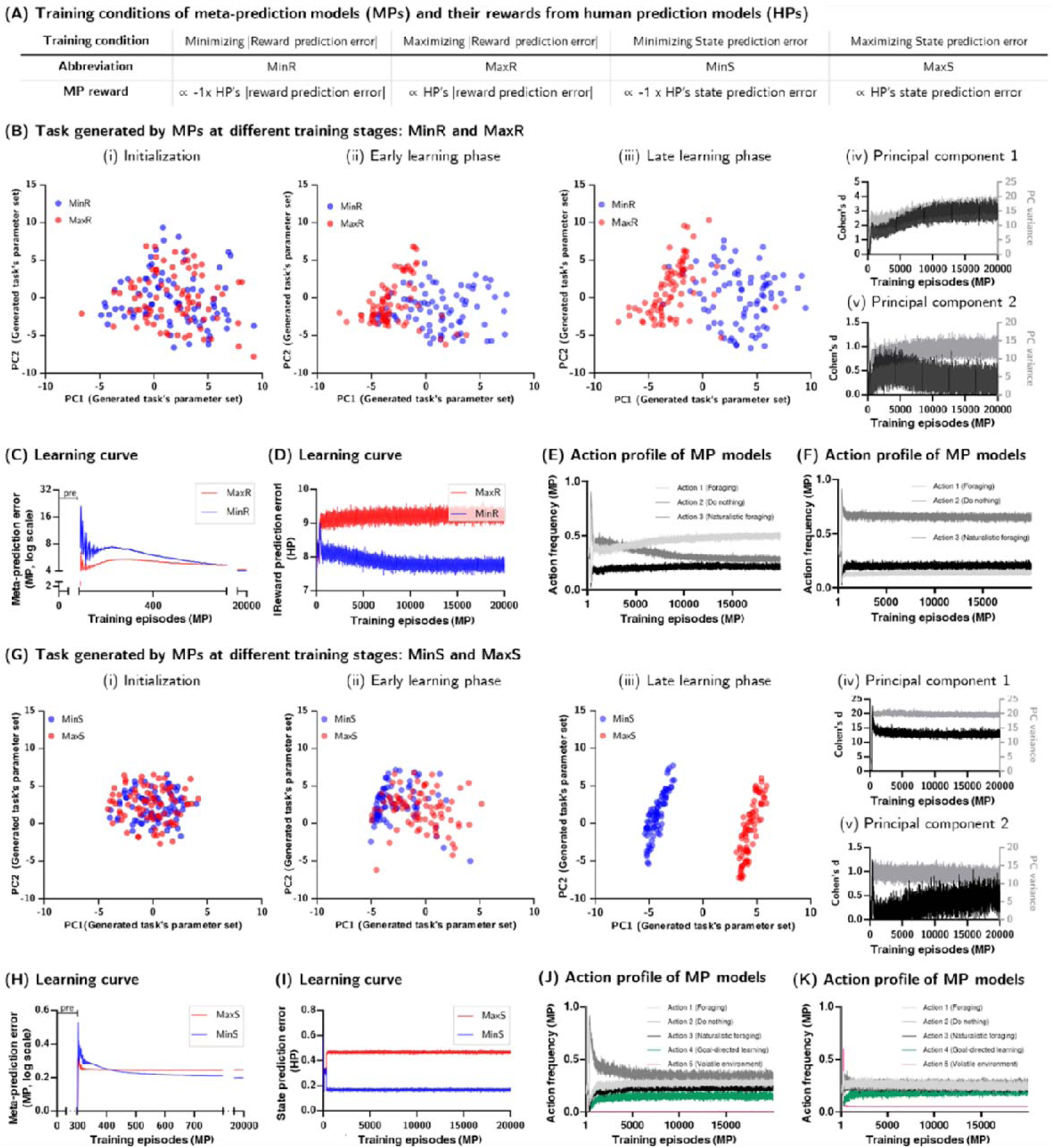
Mechanistic interpretation of meta-prediction. **(A)** Four training conditions of meta-prediction. The meta-prediction models (MP) were trained to extremize prediction errors: MinR (minimizing reward prediction error), MaxR (maximizing state prediction error), MinS (Minimizing state prediction error), MaxS (maximizing state prediction error). **(B)** Tasks generated by MPs at the initialization, early, and late training stages in MinR and MaxR. The dimension reduction analyses with PCA summarize how MPs learn to control each task parameter in two opposite training conditions. Throughout training, tasks gradually accommodate not only different training objectives (as indicated by the separability measure in (iv); Cohen’s d) but also individual variability (as indicated by the variance measure in (iv); variance). **(C)** Learning curve of the MP’s meta-prediction error. **(D)** Changes in the human prediction model (HP)’s reward prediction error during training. **(E-F)** Action profile of MPs. Shown are the changes in the MP’s action frequency, i.e., how many times the MP changed task structures. **(E)** MaxR condition**, (F)** MinR condition**. (G-K)** The same analyses as (C-F), which are for MinS and MaxS MP simulation data instead of MinR and MaxR MPs.

We conducted a total of 328 independent simulations, comprising 82 HPs’ training across 4 conditions. To mitigate potential learning biases or local minima issues in the HPs during the encoding step, we implemented a pre-training phase. This phase allowed the HPs to freely explore the task environment, enabling them to fully develop internal representations of both the task structure and reward functions. In the subsequent decoding step, MPs underwent full training while the parameters of HPs remained fixed. Despite the complexity of the training process, which involved intricate interactions between HP and MPs within each training epoch, all simulations demonstrated stable learning patterns (see Fig. 3B-K; for detailed explanation, see the ‘Training the individual MPs’ section in the Method.).

### Mechanistic interpretability of meta-prediction

Task designs generated by the meta-prediction framework help us decipher how specific environmental changes shape human prediction. Specifically, these designs serve as a direct translation of the complex dynamics underlying human reward learning into the language of environmental structure and variability. This approach enables task-level mechanistic interpretability,

For this, we examined the policy of the fully-trained MPs. The MPs for MaxR and MinR used the foraging actions only: actions 1, 2, and 3 (Fig. 2C). For maximal efficacy of the MP in controlling reward prediction error, it would be desirable for human subjects to use a model-free reinforcement learning strategy because it is the basic value learning strategy that acts on reward prediction error. To ensure this, we fixed the state-transition probability at 0.5 (highly uncertain transition) and left the goal color blank all the time (flexible goal condition; any types of rewards are achievable), the situations known to encourage model-free learning^8,9,21,22,24–26^.

The simple dimension reduction of the MP’s actions showed that the MP developed distinctly different policies to control reward prediction of the HP from a very early stage of training (Fig. 3B-(i),(ii),(iii); Cohen’s *d*-score in Fig. 3B-(iv),(v)). Moreover, the learned policies reflect individual variability, as indicated by their variance increasing with training (PC variance in Fig. 3B-(iv),(v)). The meta-prediction error of the MP converges quickly as early as 300 training episodes (Fig. 3C), effectively guiding the HP’s reward prediction error in a desired manner (Fig. 3D; MaxR > MinR, two-sample two-tailed t-test at episode 20000, n=82, *p*<0.0001).

A distinct pattern emerged in the MinR task (Fig. 3E; corresponding action sequence shown in Extended Data Fig. 1A). In the early stage, the suboptimal task favored a fixed reward function (Action 2) to stabilize the HP’s learning process. Foraging (Action 1) accelerated the reduction of the HP’s reward prediction error by narrowing the gap between actual reward values and the HP’s predictions. This preference for foraging (Action 1) persisted throughout training. Consequently, Action 1 serves to rapidly reduce the HP’s reward prediction error without compromising learnability. This strategy effectively deludes the HP into perceiving its choices as correct when they were not.

The MaxR favors a fixed environment (Action 2) with the most uncertain and complex reward distribution, in which state-transition is least predictable (*state-transition probability*=0.5) and different reward values are assigned to different outcome states (Fig. 3F; corresponding action sequence shown in Extended Data Fig. 1B). It is consistent with our intuition that uncertain and complex task structures are hard to learn.

Unlike MinR and MaxR, the MPs for MinS and MaxS aim to control state prediction error, a key variable for model-based learning. To meet this control requirement, the MPs employed all the actions, including the ones capable of changing foraging patterns, goals, and state-transition probabilities (Action 4 and 5 in Fig. 2C). For example, when goal changes, subjects must replan their behavioral policies. When reducing uncertainty in state transitions, subjects can learn about task structures rapidly. Both situations demand model-based learning strategies. To further aid highly predictable predictions of model-based learning, the initial state-transition probability was set to 0.9.

The MPs discovered distinctly different tasks for MinS and MaxS from the very early stage of training (Fig. 3G), as indicated by the rapid convergence of meta-prediction error of the MPs (Fig. 3H). As a result, it effectively guides HP’s state prediction error (Fig. 3I; MaxS > MinS, two-sample two-tailed t-test at episode 20000, n=82, *p*<0.0001). In the MinS (Fig. 3J; corresponding action sequence shown in Extended Data Fig. 1C), the MP maintained its state-transition probability to match the current predictions of the HP (avoiding Action5). In the MaxS, on the other hand, the MP used most of the actions (Action1-4) almost equally to create inconsistent reward situations (see Fig. 3K) except Action5. Here the MP changes state transitions to the least predictable condition (by taking a one-time Action5 to change *state-transition probability* from 0.9 to 0.5) in the beginning of trials and keeps it all times (corresponding action sequence shown in Extended Data Fig. 1D). The ablation analyses also confirmed the significant contribution of the MP’s key actions (Action1 for MinR, Action 2 for MaxR, and Action5 for MinS and MaxS, see Extended Data Fig. 2).

### Validation of meta-prediction through cross-subject shuffle test

To confirm that the MP effect generalizes to new human subjects who were not part of the initial cohort used to develop the HP and MP, we first sought to identify the most generalizable *meta-prediction* model (MP) from our simulation results (Fig. 3). We employed a MP-HP shuffle test, inspired by cross-validation in machine learning (See Methods for more details). This test evaluates each MP’s average performance across all the other HPs not used during training, estimating out-of-sample performance.

Notably, across all four training conditions (MinR, MaxR, MinS, MaxS), the shuffle test identified subject-independent MPs that consistently performed well across all 82 HPs within each condition (Fig. 4A). These MPs demonstrated performance comparable to subject-specific MPs in all conditions (paired two-tailed t-test with Bonferroni correction for multiple comparison, *p*>0.999 (MinR, n=82; MaxR, n=82; MaxS, n=82), p=0.9790 (MinS, n=82), detailed subject-independent MP policies are provided in Extended Data Fig. 3). These findings suggest that, for these four key conditions, a single representative MP policy may generalize across individuals, without the need for individual-specific task adaption or calibration.

**Fig 4.**
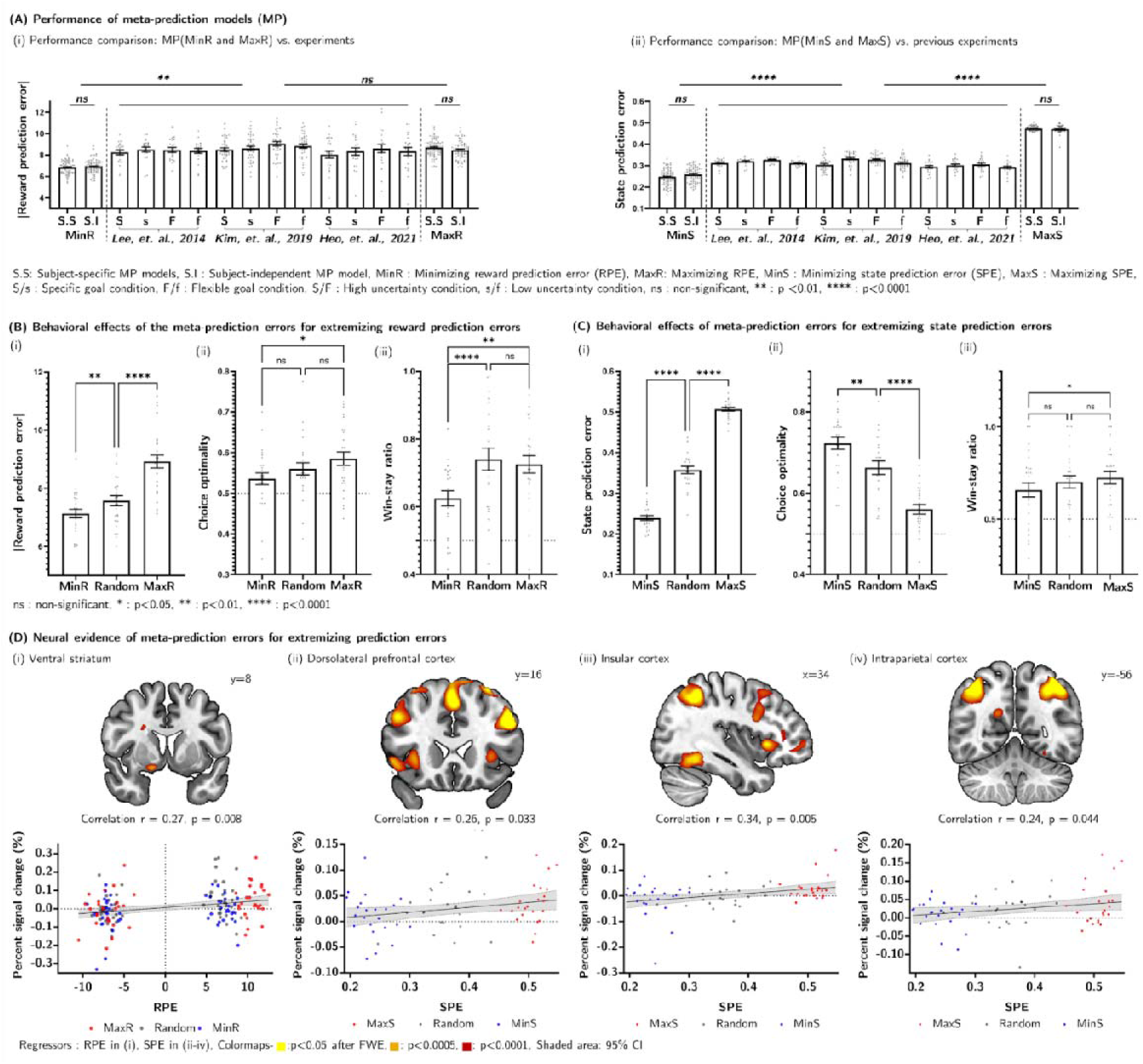
Behavioral and neural effects of meta-prediction. **(A)** Performance of meta-prediction models (MP) was evaluated based on the prediction errors generated by human prediction models (HP). Performance comparisons were made with the previous studies^8,21,22^. Prediction error was assessed under four training conditions: MinR, MaxR, MinS, and MaxS (See Fig. 3A). S.S denotes subject-specific MP, trained individually on each HP. S.I. denotes subject-independent MP, a single MP identified through the HP-MP shuffle test that achieved the best average performance across all HPs. We validated the generalizability of S.I. by demonstrating that its performance was comparable to that of its subject-specific counterparts, S.S. We then used S.I. for the rest of the analyses. In addition to this direct performance, we also confirmed that the generated task designs elicit greater variability in human prediction error (Fig. 4A-MP reward), compared to the prior two-step reward learning tasks^8,21,22^, which relied on the hand-crafted environments. **(B)** Behavioral effects of the MPs in MinR and MaxR. The effect was measured based on (i) the absolute reward prediction errors, estimated from the model-free reinforcement learning (RL) fitted to individual subjects’ behavior, (ii) the choice optimality, and (iii) the win-stay ratio of individual subjects. **(C)** Behavioral effects of the MPs in MinS and MaxS. The state prediction errors were estimated from the model-based RLs fitted to individual subjects’ behavior. **(D)** Neural evidence of MPs. Activities in the ventral striatum^2,7,8^ correlated with reward prediction errors (i), while activities in the lateral prefrontal cortex^7,8^(ii), insula^8^(iii), and intraparietal sulcus^7,8^ (iv) correlated with state prediction errors. The percent signal change analysis was performed on a 5mm radius. Error bars represented SEM across subjects. Asterisks (*) indicate significance levels: * p<0.05, ** p<0.01, *** p<0.001 (two-tailed paired t-test with n = 82 for (A), n=26 for (B)-i, (C)-i, (C)-iii, n=23 for (B)-ii, (C)-ii, (C)-iv). Clusters showing significant effects at a threshold of p<0.001 (uncorrected) with a minimum cluster size of 100 voxels are displayed in yellow in the brain images located in the upper row of the sub-figure (D).

Moreover, both subject-specific and subject-independent MPs outperformed previous experimental designs in separating state and reward prediction errors - particularly those intended to dissociate model-based and model-free learning systems^8,21,22^. In the MinR condition, MPs tailored to minimize reward prediction error reduced the HPs’ reward prediction errors beyond those observed in prior experiments, while in the MaxR condition, they increased reward prediction errors to a comparable degree (Fig. 4A-i; paired two-tailed t-test with Bonferroni correction for multiple comparison, p<0.013 (MinR vs previous experiments, n=82), *p*>0.999 (MaxR vs previous experiments, n=82)). Similarly, the MPs designed to extremize state prediction error yielded better performance in both MinS and MaxS compared to the prior experiments. (Fig. 4A-ii, paired two-tailed t-test with Bonferroni correction for multiple comparison, p<0.001 (MinS vs previous experiments, n=82; MaxS vs previous experiments, n=82)). These findings demonstrated the ability of our MP framework to generate calibrated task environments that effectively modulate human prediction errors.

### Behavioral and neural effects on human reward learning

Using the task generated by the subject-independent MP, we conducted two fMRI experiments with 49 human subjects in total. In the first experiment, 26 participants performed the reward prediction error tasks (MinR and MaxR), and in the second experiment, 23 participants completed the state prediction error tasks (MinS and MaxS). Subjects learned the task well in all cases, as evidenced by the above-chance-level choice optimality (one-sample two-tailed t-test, *p*=0.022 (MinR, n=26); *p*=0.004 (MaxR, n=26), *p*<0.0001 (MinS, n=23; MaxS, n=23)). The choice optimality was defined as the proportion of a subject’s choice that matches those of an optimal model-based reinforcement learning algorithm with complete task information, including reward function and state-transition probability.

As an initial validation, we tested whether the subject-independent MPs modulated subjects’ prediction errors. The reward and state prediction errors were quantified by fitting model-free and model-based reinforcement learning algorithms to individual choice behavior, respectively. In the first experiment (MinR and MaxR), we found that the MP effectively extremized reward prediction errors (MaxR > MinR in Fig. 4B-(i), paired t-test, p<0.0001, n=26). The second experiment (MinS and MaxS) also confirmed that the MP effectively extremized state prediction errors (MaxS > MinS in Fig. 4B-(ii), paired t-test, p<0.0001, n=23).

We then examined the influence of MPs on the reward-related behavior of human subjects. The MPs for prediction error minimization and maximization resulted in statistically significantly different choice behavior patterns. Choice optimality, a standard behavioral index for goal-directed learning^23^, was higher in the MaxR condition compared to the MinR (Fig. 4C-(i); paired t-test, p=0.0145, n=23), while it was higher in the MinS than the MaxS (Fig. 4C-(iii); paired t-test, p < 0.0001, n = 26). The win-stay ratio, a typical behavioral metric for habitual learning^27^, was higher in the prediction error maximization (MaxS and MaxR) compared to the minimization (MinS and MinR) (Fig. 4C-(ii); paired t-test, p=0.0141, n=26; Fig. 4C-(iv), paired t-test, p=0.004, n=23). This measure was defined as the proportion of a subject’s first-stage action choices that replicated those from the preceding trial, conditional on that trial yielding a high-valued reward (for details, see the ‘Win-stay ratio’ section in the Method).

Lastly, we evaluated the neural effect of MP on reward learning using model-based fMRI analyses (see the fMRI acquisition and fMRI analysis section in Method). The whole-brain analyses confirmed the previous findings^2,7,8^ that the reward prediction error was reflected in the activity patterns of the ventral striatum (Fig. 4D-(i)), while the state prediction error was encoded in the neural activity of multiple cortical areas, including dorsolateral prefrontal cortex, insula, and intraparietal sulcus (Fig. 4D-(ii),(iii),(iv)), fully consistent with the previous studies^7,8^. We also examined specific effects of MP in the core area known to guide reward learning. For this, the neural activity was extracted from the BOLD signals in the identified cortical regions and the ventral striatum (for detailed ROI definitions, see the ‘percent signal change’ section in Methods). The neural effect of MP in these areas survived after the most stringent Bonferroni correction for multiple comparisons. The neural activity change was significantly greater in MaxR/MaxS compared to MinR/MinS (Percent signal change plots in the bottom of Fig. 4D).

### Decoding individual goal-habit bias through meta-prediction

The demonstrated efficacy of the meta-prediction framework in modulating human reward learning, both behaviorally and neurally, motivated us to further investigate its potential for quantifying innate reward learning biases, particularly the dynamic balance between goal-directed and habitual learning strategies^3,11,28–33^. This goal–habit bias represents a critical dimension of cognitive flexibility, whose dysregulation has been implicated in psychiatric disorders such as addiction or obsessive compulsive disorder^34–40^. To assess whether our meta-prediction framework could capture such bias, we trained MPs to modulate the key variable known to guide transitions between goal-directed and habitual control^8,21,22,41,42^: long-term prediction errors of the HPs (Fig. 5A). They are defined as the reward and state prediction errors over multiple trials. For consistency, we used the same simulation settings as before (Figs. 3 and 4). The HPs used in our simulation were previously fitted to data from 82 human subjects, thus serving as proxies for individual human reward learning.

**Fig 5.**
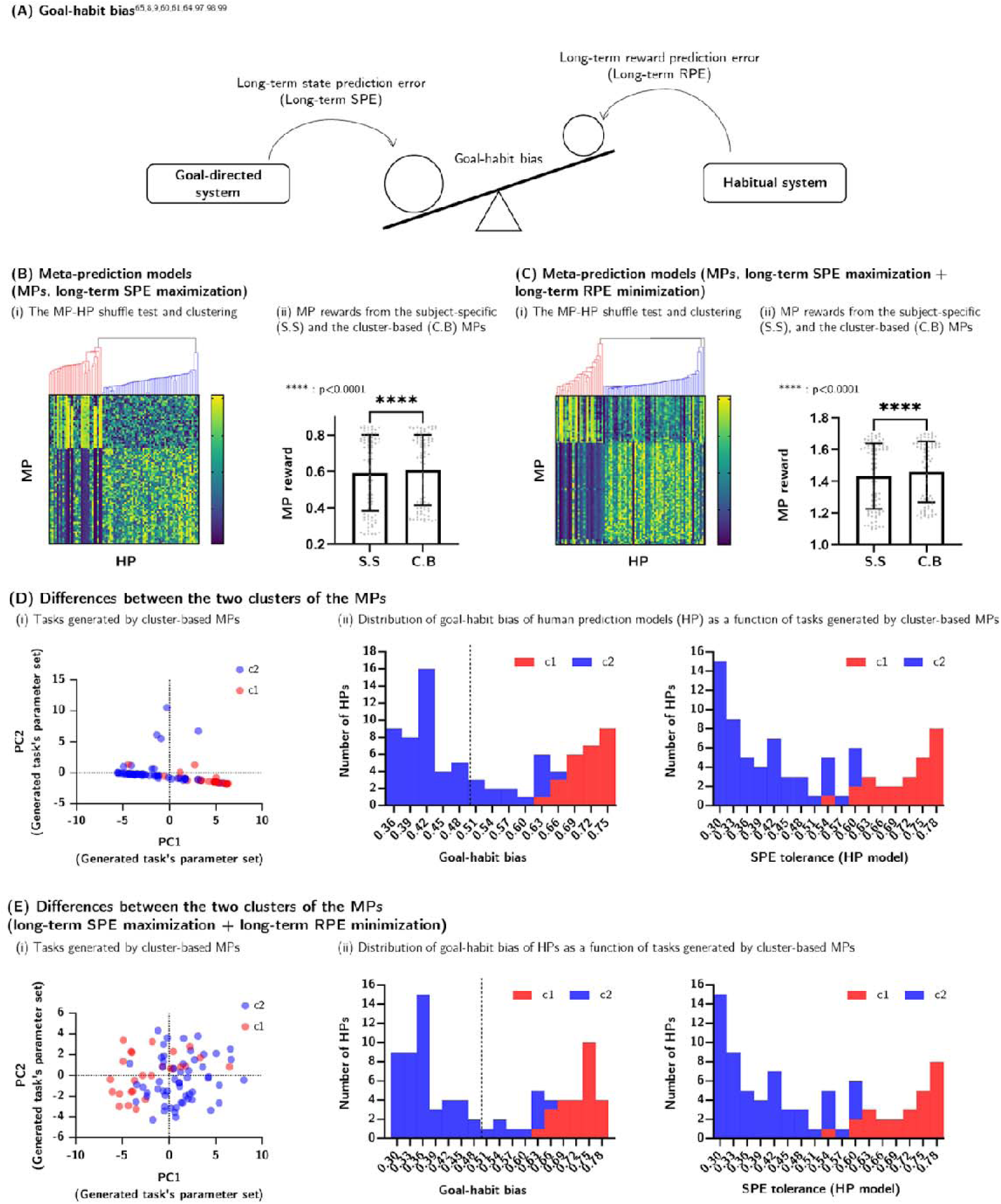
Individual goal-habit bias classification with meta-prediction. **(A)** Effect of long-term prediction error (PE) on goal-directed reinforcement learning (RL) adopted from *Yoshida et al.(2014)*^41^. Earlier works showed that the reward/state PE (RPE/SPE) over multiple trials (long-term PE) is the key variable to explain bias towards two different reward learning strategies: goal-directed and habitual learning^8,21,41,42^. This view suggests that a meta-prediction model (MP) to control long-term PE could decode the goal-habit bias. **(B)** MPs trained for long-term SPE maximization. (i) MP-human prediction model (HP) shuffle test. We trained MPs for long-term SPE maximization (long-term MaxS). Shown are the MP rewards obtained from the frozen MPs applied to each HP. Two MP clusters were found, as indicated by c1 and c2. (ii) In terms of MP rewards, the cluster-based subject-independent MPs (C.B.) are better than a single subject-independent MP (S.I.) and multiple subject-specific MPs (S.S). (Wilcoxon test with Bonferroni test for multiple comparison; n=82, *p*<0.0001 for both S.S vs C.B and S.I vs C.B). **(C)** MPs were trained for long-term SPE maximization + long-term RPE minimization. The same analyses as (B), except for MPs’ target, which was the long-term SPE maximization + long-term RPE minimization (dual objective long-term MaxS + MinR, Wilcoxon test with Bonferroni test for multiple comparison; n=82, *p*<0.0001 for both S.S vs C.B and S.I vs C.B) **(D)** (i) MP-generated tasks for long-term SPE maximization (dimension reduction analyses). Two different types of MP-generated tasks were indicated by red (c1) and blue (c2) clusters. c1 and c2 were labeled from the (B). (ii) Distribution of goal-habit bias (left) and SPE tolerance (right) of HPs was color-coded by the two clusters found in (C). The goal-habit bias was measured as an innate preference to habitual RL (w in *Daw et al.(2011)*^10^; P_mb_ in *Lee et al.(2014)*^8^). The SPE tolerance is the key parameter to reflect the sensitivity to its own PE^8^. **(E)** The same analyses as (D), except for MPs for long-term SPE maximization + long-term RPE minimization. c1 and c2 were labeled from the (C). Although (D) and (E) were based on independent experiments with independently performed clustering, the SPE tolerance parameter histograms were identical, as both experiments resulted the same HP clusters (see Extended Figure 8).

To identify an optimal training regime, we trained MPs under four training conditions: long-term MinR, MaxR, MinS, and MaxS. As predicted by the earlier study on the effect of long-term prediction error on goal-habit bias^8^, the MP trained under the long-term MinR condition facilitated goal-to-habit transition by providing a stable and predictable environment (Extended Data Fig. 5B. In contrast, the MP under the long-term MaxS condition effectively disrupted the goal-habit bias by placing high demands on goal-directed learning (Fig. 5B, Extended Data Fig. 5C). Building on this insight, we introduced a combined training regime, long-term MaxS + MinR, to accelerate the transition from goal-directed to habitual control (Fig. 5C).

The trained MPs enabled rapid transitions from goal-directed to habitual control in a subject-specific manner (Extended Data Fig. 6H). Consequently, any decline in MP performance on different HPs is likely to reflect individual variation in goal–habit bias. To probe this, we performed an MP-HP shuffle test, evaluating each MP’s performance across all MP-HP combinations based on the total reward accrued by the MP. This analysis revealed two distinct MP-HP clusters, wherein one MP group effectively extremized the long-term prediction errors of one subset of HPs, but not for the other (labeled c1 and c2 in Fig. 5B,C-(i), detailed clustering results in Extended Data Fig. 7H). Notably, we identified a subject-independent MP within each cluster, which exhibited the best average performance across all HPs, surpassing both the subject-independent and subject-specific MPs from the earlier simulations of Fig. 4 (Fig. 5B,C-(ii), Wilcoxon test, p<0.0001 (long-term MaxS, n=82; long-term MaxS+MaxR, n=82)).

Interestingly, similar clustering patterns were found under more complex training regimes such as long-term MaxS+MaxR and long-term MaxS+MinR (see Extended Data Fig. 7E,H), with consistent grouping patterns (see Extended Data Fig. 8). We also found that these MPs tend to favor action repetition policies consistent with those derived from simpler training conditions, such as long-term MaxS, suggesting underlying structural inductive bias in task design (see Extended Data Fig.9). This highlights an intriguing possibility for compositional task design. For instance, complex tasks can be constructed by combining a small set of primitive MPs trained under foundational conditions (MinR, MaxR, MinS, and MaxS).

Building on the above findings, we evaluated whether these clusters (c1 and c2) are associated with HPs’ goal-habit bias. This bias was defined as the extent to which a model-based reinforcement learning algorithm better accounts for each subject’s behavior compared to a model-free approach, quantified using likelihood values. Interestingly, the HP’s goal-habit bias showed biomodal distributions that closely aligned with the two clusters identified in the MP-HP shuffle test (Fig. 5D,E-(i)). Note that this cluster-level alignment was found solely from MP performance, without access to any HP-specific information. This presents a new possibility for decoding individual goal-habit bias. By either assessing MP performance or simply aggregating the prediction error from a new human subject, it may be possible to infer not only an individual’s goal-habit bias (Fig. 5D,E-(ii)) but also a key parameter underpinning this bias: state prediction error tolerance (Fig. 5D,E-(ii)).

## Discussion

This study addresses a fundamental challenge in reward learning task design: overly stable environments constrain learning opportunities, and highly uncertain ones render reward learning intractable. To address this dilemma, we proposed a novel conceptual framework, *meta-prediction*, which integrates two Bellman equations: one that emulates human reward learning, while the other learns to generate task design by predicting the first. This approach offers three key advantages. First, it enables optimal task design by navigating the tension between simple and uncertain environments. Second, It provides mechanistic interpretability, translating the complex nature of human reward learning into task design. Third, it supports compositional task design, revealing how complex tasks can be assembled from primitive, model-driven task components generated by our framework.

While traditional data-driven approaches for task optimization or task adaptation primarily rely on observable behavior, our framework targets a latent variable underlying human reward learning: prediction error. This approach enables a deeper understanding of how internal expectations shape external behavior. Specifically, we demonstrated that modulating two distinct types of prediction errors - reward-based and state-action-state transition-based - yields distinctly diverse changes at the behavior and neural levels. Behaviorally, these manipulations could influence motivation^44^ and difficulty^45^, and neurally, they engage distinct pathways, including the basal ganglia^28,29,33^ and prefrontal cortex^39,32^.

Beyond observable choice behavior, our framework provides a window into the innate biases underpinning human reward learning. Specifically, we demonstrated that the MP performance profile, derived from the MP-HP shuffle test, revealed two latent dimensions: the degree of goal-directed behavior (“goal-habit bias” in Fig. 5D,E-(ii)) and its key variable to guide the behavioral transition from goal-directed to habitual learning^8,22^ (“SPE tolerance” in Fig. 5D,E-(ii)). This allows for the readout of the individual bias of goal-directed learning without model-fitting, which can be time-consuming and prone to interpretive pitfalls. Instead, our meta-prediction offers a compositionally scalable, mechanistically interpretable, and personalized means to optimizing task design (Extended Data Fig.9).

Previous researches have demonstrated the potential of using computational cognitive models to generate tasks that influence human decision-making^16,17^. However, these approaches have limited scalability and applicability due to their dependence on a narrow task parameterization scheme (Single-step Markov decision task^16^, Go-NoGo test^17^) and the repeated use of near-identical task conditions (Multiround trust task^17^). Moreover, these models primarily target explicit human behaviors (left/right choices^16,17^, amount of payback^17^), while overlooking the underlying learning strategies underpinning such behaviors. Although these models are trained on observable human actions, they provide limited guarantees regarding alignment between the decision-making processes of humans and artificial neural networks (ANNs). A recent study^46^ suggested a method for value alignment between humans and ANNs using the hidden cognitive variables, such as causal uncertainty. Nonetheless, it does not yet establish a direct correspondence with the underlying neural substrates guiding human reward learning processes.

Our framework addresses this limitation through a novel approach: the Meta-Prediction model (MP) generates or optimizes parameterized tasks, while the Human Prediction model (HP) provides predictions within these tasks. We have demonstrated that MPs successfully generate tasks across various training conditions, even when their task space differs from that of the HPs. Future work could benefit from our HP-MP framework by generalizing it to an encoder-decoder architecture, with HP serving as encoders and MP as decoders. This architecture is interpretable, as the encoder provides a direct estimation of human prediction error, while the decoder offers interpretation at the task level. This encoder, which can be exchanged with diverse structures of HPs^47,48^, also offers generalizability and unified insights into human behavior prediction and associated task generation

Our meta-prediction framework has wide-ranging applications across various domains. One key area is curriculum learning^49^, where our framework enhances task sequencing by streamlining a progression of learning experiences that promote goal-directed behavior without compromising overall learnability. In decision neuroscience, it helps fine-tune existing task structures for better dissociation between state and reward prediction errors. In smart education, it can enhance learning efficiency while accommodating individual variability^50^. Additionally, our framework can guide the diagnosis and treatment of mental disorders associated with learning and prediction. By leveraging models of abnormal behavior distributions in various conditions, the meta-prediction module could generate tailored tasks to facilitate diagnosis or intervene in the latent processes for cognitive behavior therapy. In summary, our meta-prediction theory offers a fundamental solution to value alignment between humans and AI models. Training a meta-prediction module to minimize prediction error can be evolved into the process of aligning the relevant value functions^51^ of human subjects with those of an arbitrary AI model, implying the AI model’s learning of the human learning process, or the ‘meta-learning’ AI model.

## Materials and Methods

### General Stimuli

We used fractal images to represent the initial state, four intermediate states following the player’s first action, and four terminal states associated with rewards. Each subject observed nine fractal images randomly selected from a set of 126 images at the beginning of the task. A square box was displayed beneath the state image to indicate the target goal color (red, blue, yellow, or white). In the terminal state (i.e., the final phase of each trial), a state-associated coin (red, blue, yellow, or gray) and its value appeared below the target goal color image. Coin acquisition depended on matching the target goal color, which was displayed alongside the coin value. The gray coin consistently had a reward value of zero throughout the task, while the values of the other colored coins were determined at the start of each experiment. The ranking and cumulative scores were presented below the coin value to maintain participant engagement.

### General Tasks

The experimental setup for fitting the human prediction model was adapted from previous two-stage Markov decision tasks^7,8^. We then modified this setup for meta-prediction model training and its application to novel human subjects. The basic structure of the experiment was the same for both simulation and real-human subject tasks. Participants pressed one of two buttons (‘Right’ or ‘Left’) based on the current state image and goal color. If no choice was made within 4 seconds of the state image presentation, the computer randomly selected an action to continue the task. The next state appeared between one and four seconds after the action. Following the final state and reward presentation, a new game began two seconds later, starting with the same initial state image.

Each action led participants to one of 2 possible next states based on state transition probabilities that the participants had to infer implicitly. The meta-prediction model set the state transition probabilities of either [0.5, 0.5] or [0.9, 0.1]. The state-action-reward mapping remained constant for each participant across all sessions. Participants learned this mapping and inferred the current state transition probabilities during a pre-training session, which was not included in the performance evaluation. The participants then applied their knowledge during five main sessions. Each session consisted of 80 game trials, organized into four blocks of 20 trials.

Prior to the pre-training session, participants were instructed about potential changes in state transition probabilities and the implications of the goal color image. However, they were not informed of the specific values of the colored coins or the exact state images they would encounter. The colored coins, except for gray, represented ‘high,’ ‘medium,’ and ‘low’ reward values – initialized at 40, 20, and 10 rewards, respectively – and these values remained fixed as upper boundaries throughout the task. During simulation, rewards for non-target goal coins were deactivated and hidden under specific goal conditions until reactivation by the meta-prediction models.

At the start of each game, participants could identify the current goal color from the displayed goal color indicating images. In the flexible goal condition (white box), participants could obtain coins at terminal states regardless of color. In the specific goal condition (red, yellow, or blue), coin acquisition was restricted to the trials where the coin color matched the target goal color. These rules were explained to the participants before the pre-training session.

We integrated foraging paradigms into the conventional two-stage Markov Decision Task to incentivize explorative behaviors and mitigate habitual decision-making strategies. The initial value of the coins was set as the upper boundary threshold, with reward values below two decimal points systematically eliminated. To promote foraging behavior and discourage habitual actions, the value of the acquired reward coin was devalued by 65%-85%.

Each block of tasks was administered according to either (i) the task environments generated by the meta-prediction model or (ii) a pseudo-random policy where action selections followed a stochastic distribution with controlled frequencies to maintain outcome equilibrium. After completing each block, participants were asked to rate task difficulty on a 5-point scale. This subjective rating, however, was not incorporated into the meta-prediction model.

### Human prediction models (HPs)

We used the same arbitration model of reinforcement learning (RL) described in the previous paper^8^. The model consisted of three primary components: a model-free learner, a model-based learner, and an arbitrator.

The model-free learner utilized the SARSA algorithm^1^, a standard approach in RL. The model-based learner employed forward learning^7^ for state transition updates and backward planning to address predictable changes in task structures. Both learners generated distinct prediction errors following each action: the model-free learner produced reward prediction error (RPE), while the model-based learner generated state prediction error (SPE). The arbitrator evaluated each learner’s reliability based on the history of prediction errors and modulated their relative contributions through a dynamic equilibrium mechanism^8^. Furthermore, the arbitrator computed the integrated Q-action values from both learners and selected actions via the soft-max function^7,50^. For a detailed algorithm description, see the supplementary data S5 (HP algorithm).

The HPs were trained on behavioral data collected from 82 participants across two experiments: 22 adults (6 females, mean age 28, age range 19-40) from the USA and 60 adults (31 females, mean age 22.75, age range 19-35) from South Korea. The same two-stage Markov decision task, excluding the foraging condition, was used for both experiments. Both studies received ethical approval from their respective Institutional Review Boards – Caltech for the U.S. experiment and KAIST for the South Korean experiment.

### Meta-prediction model (MPs)

The MPs chose one from five distinct, pre-defined options (see Figure 2C). The first three options were available to both SPE and RPE control, which focused on restoring or maintaining the value of the devalued coins: (i) *Foraging* – kept the task structure unchanged after reward devaluation, (ii) *Stable environment* – targeted previously devalued coins of the same color, and (iii) *Naturalistic foraging* – targeted coins of the different colors other than those previously devalued. These three options above increased the targeted coin values by multipliers between 1.18 and 1.54, with values capped at a predetermined maximum. Next, the additional two options were exclusive to SPE control: (4) *Goal-directed learning* – alternated between specific goals (which randomly selected colors at the beginning of each game) and flexible goals, and (5) *Volatile environment* – modified the state transition probabilities between (0.9, 0.1) and (0.5, 0.5).

We implemented the MPs using the double-DQN algorithm^19,20^. The MPs featured a fully connected architecture with a single hidden layer containing 64 neurons. The input vector consisted of 20 dimensions: 16 dimensions for the rewards at terminal states, two dimensions for transition probability, and two dimensions representing SPE and RPE values for the current agent. The action selection of the MPs was guided by an epsilon-greedy policy (epsilon = 0.1) to ensure a balance between exploration and exploitation. The models were trained using the RMSprop algorithm^51^ with mean squared error as a loss function. The replay buffer had a capacity of 4500 samples. The training was conducted with a batch size of 32. Each MP was trained for 20,000 episodes, which included a pre-training phase.

The reward functions for the MPs varied depending on their training conditions. When an MP was designed to maximize a single immediate or long-term prediction error, it received a reward directly proportional to that target prediction error. Conversely, MPs designed to minimize a single target received a reward inversely proportional to the corresponding prediction error.

For models trained to control multiple target prediction errors, the reward function was defined as a combination of the individual reward functions associated with each target. For example, an MP trained to maximize long-term state prediction error while minimizing long-term reward prediction error received a reward equal to the sum of two sub-rewards: one directly proportional to the long-term state prediction error, and the other inversely proportional to the long-term reward prediction error.

The MP’s target prediction errors included the HP’s absolute reward prediction error, state prediction error, long-term reward prediction error, and long-term state prediction error.

### Training the individual MPs

We implemented the pre-training stage for both the HP and MPs to replicate the experimental setting of the original human subject experiments. During the first 100 episodes, the HPs explored the default foraging task by performing random actions. For the MPs, we first detached them from the task for the initial 25 episodes out of 100 episodes. Afterwards, we had them perform random actions for the remaining 75 episodes to explore their action space. For algorithm description, see the supplementary data S4 (HP pre-training) and S6 (MP training).

### Analysis of the behavioral dynamics of tasks generated by MPs during training

In relation to Figure 2B, we conducted a comprehensive analysis on the behavioral characteristics of tasks generated by the MPs, focusing on their dynamics across the initialization, early, and late training stages. We achieved it by systematically evaluating task parameters at each stage of training.

First, we captured snapshots of the tasks generated by the MPs during each training episode. Each snapshot consisted of a sequence of trials within the episode and the corresponding task parameters – transition probability and coin reward values – manipulated by the MP.

Next, we categorized the collection of snapshots into two distinct groups after vectorizing the tasks. Each task was encoded as a vector, with episodes comprising 20 trials represented by 80-dimensional vectors (20 trials × 4 features: one feature for transition probability and three for coin rewards). These vectors were grouped into two categories based on the MP’s target prediction errors.

We then performed principal component analysis (PCA) on the tasks in each group, extracting the two principal components with the highest variance. Finally, we calculated Cohen’s distance^52^ between these components, aligning the measurement with the direction of the MP’s prediction error control.

### Ablation study to evaluate the contribution of the MPs’ key action

To highlight the significance of the MP’s key action, we conducted a systematic ablation study by strategically reducing the action space while preserving the consistent experimental setup for the MPs’ training. In these control tasks, MPs incorporating SPE were limited to four actions (i.e., one action was removed from the original five). In contrast, those employing RPE were restricted to two actions (i.e., one action was removed from the original three). Importantly, all other experimental settings remained identical to the original MP training framework, ensuring a controlled comparison. This approach enabled us to examine the sensitivity of the models to changes in action space and to assess their controllability across varying control task objectives (e.g., MinR, MaxR, MinS, MaxS).

### Acquisition of the generalized MPs

We systematically collected subject-specific task environments generated by trained MPs, each of which corresponded to a distinct HP. A comprehensive set of 82 task environments was generated for each MP objective. These meticulously constructed task environments were subsequently applied across all HPs, resulting in 82 × 82 sets of behavioral data, wherein HPs responded to the task environments generated by the MPs. For each HP-MP pair, we conducted a simulation of 100 episodes and computed the average reward achieved by the MP across these episodes.

The MP that generated the task environment demonstrating the highest average reward across all HPs was identified as the subject-independent MP. To rigorously assess performance, we conducted a paired *t*-test to statistically compare the rewards obtained by the subject-independent MP against those generated by individual HP-specific MPs.

### Participants

A total of 26 right-handed volunteers (15 females, mean age 22.5, age range 19-35) participated in the RPE control task, while 23 right-handed volunteers (9 females, mean age 24, age range 19-31) took part in the SPE control task, both of which included fMRI scanning. Participants were pre-screened to exclude individuals with a history of neurological or psychiatric illnesses. All participants provided informed consent, and the study was approved by the KAIST Institutional Review Board.

### Behavioral measures

#### Choice optimality

The choice optimality of individual participants evaluates whether they had correctly learned the task structure and adapted their behavior across different task blocks. It is defined as the proportion of optimal first-stage actions relative to the total number of first-stage choices. An action is considered optimal if it has the highest expected value, assuming full knowledge of both the coin reward values and the state transition probabilities.

#### Win-stay ratio

The win-stay ratio estimates individual habitual learning bias, which reflects the tendency to persist with previous action choices that led to rewarding outcomes. Rewarding outcomes are defined as coin rewards that exceed the average value of the current coin set. For example, if the coin values were (Red: 35, Blue: 15, Yellow: 7, Gray: 0), the rewarding outcomes would be the red and blue coins, as their values exceed the average (14.25). The win-stay ratio is then calculated as the proportion of trials in which the first-stage action is repeated following a trial that yielded a rewarding outcome.

#### Reaction time

We analyzed the effect of task control conditions on reaction time (RT) in both RPE and SPE control tasks. In both conditions, there were no significant differences between RT values. (RT MinR = 0.68s; RT MaxR = 0.71s, RT MinR vs MaxR paired *t*-test; *p*=0.174; RT minS = 0.72s; RT maxR = 0.74s, RT MinR vs MaxR paired *t*-test; *p*=0.58, see Extended Data Figure 4 for details.)

### fMRI data acquisition

Functional imaging for the RPE task was conducted using a 3T Philips Achieva scanner located at the Korea Basic Science Institute (KBSI). In comparison, the SPE task was performed on a 3T Siemens Verio scanner located at the KAIST fMRI center. Both scanners used 32-channel radio frequency coils. High-resolution structural images were acquired using MPRAGE sequences, with a resolution of 1.2 mm × 1.2 mm × 1.2 mm for the RPE task and 1 mm × 1 mm × 1 mm for the SPE task.

Functional scans were acquired using a one-shot echo-planar imaging sequence (RPE: TR = 2500 ms; TE = 35 ms; FOV = 100 mm; flip angle = 79°; SPE: TR = 2500 ms; TE = 30 ms; FOV = 100 mm; flip angle = 90°). Both tasks acquired 45 slices, with voxel resolutions of 3.5 mm × 3.5 mm × 3.5 mm for the RPE task and 3 mm × 3 mm × 3 mm for the SPE task.

### fMRI Data analysis

We analyzed the fMRI data using the SPM12 software package (Wellcome Department of Imaging Neuroscience, Institute of Neurology, London). Slice-timing correction was applied to the functional images to address differences in acquisition times across individual slices within each scan. The images were then motion-corrected, spatially normalized to a standard echo-planar imaging (EPI) template to account for anatomical variability between participants, and smoothed using a 3D Gaussian kernel with an 8 mm full width at half maximum (FWHM). The processed dataset was subsequently subjected to statistical analysis.

### GLM analysis

We employed a General Linear Model (GLM) to perform whole-brain analyses of human decision-making processes. The model included regressors for stimulus onsets (R1), parametric regressors encoding target prediction errors (R2-RPE or SPE) at the same time points, and the movement-related regressors (R3-R8). For each GLM run at the single-subject level, orthogonalization of the regressors was disabled to preserve the correlated information between SPE and RPE. A whole-brain FWE correction (p < 0.05) was applied to account for multiple comparisons. A high-pass filter (cutoff = 129 s) was used to remove low-frequency noise from the data. All reported brain coordinates were based on MNI coordinates.

### Percent signal change analyses

We selected spherical regions of interest (ROIs) based on the GLM results to examine prediction errors. The SPE ROIs were defined as 20 mm radius spheres centered at (30, −56, 44), (48, 16, 46), and (34, 22, 0) for the intraparietal sulcus, dorsolateral prefrontal cortex, and insular cortex, respectively. The RPE ROI was defined as a 5 mm radius sphere centered at (−14, 8, −12) for the ventral striatum. All brain coordinates were based on MNI coordinates. The difference in sphere radius was chosen based on the varying sizes of the relevant brain regions.

We performed a percent signal change analysis using the rfxplot toolbox (http://rfxplot.sourceforge.net/) to investigate prediction error onsets across different MP objectives (minimizing, maximizing, and pseudo-random). Due to the relatively slow block frequency (20 trials per block ≈ 15 minutes per block), we turned off high-pass filters during the 1st level GLM analysis prior to conducting the percent signal change analysis.

### HP-MP shuffle analysis

After acquisition of the HP-specific MPs for controlling long-term prediction errors, we assessed the controller reward ranking for each MP within each HP. Each MP was assigned an 82-length vector representing its ranking across specific HPs. We performed hierarchical clustering based on the Euclidean distance between the controller reward ranking vectors.

Task environments that did not involve maximizing long-term state prediction errors showed no clear separations (Extended Data Fig. 7-A,B,D,F,G). In contrast, three task conditions that involved maximizing long-term state prediction errors exhibited similar cluster separations (Extended Data Fig. 7-C,E,H), differing by only about 2-3 subjects (Extended Data Fig. 8).

For further cluster analysis, we selected clusters that maximized long-term state prediction errors + minimized long-term reward prediction errors. The same method used for prediction error analysis was applied to the principal component analysis of the task environments.

## Data availability

The raw behavioral data and fMRI results are available for download at https://github.com/brain-machine-intelligence/meta_prediction_2024.

## Code availability

The simulation codes are also available for download at https://github.com/brain-machine-intelligence/meta_prediction_2024.

## Acknowledgements

This work was funded by the National Research Foundation of Korea(NRF) grant funded by the Korea government(MSIT).(RS-2024-00341805), Institute of Information & communications Technology Planning & Evaluation (IITP) grant funded by the Korea government (MSIT) (No. RS-2023-00233251, System3 reinforcement learning with high-level brain functions), the National Research Foundation of Korea(NRF) funded by the Korean government (MSIT) (No. RS-2024-00439903), Institute for Information & communications Technology Promotion(IITP) grant funded by the Korea government(MSIT)(No.RS-2019-II190075 Artificial Intelligence Graduate School Program(KAIST), Electronics and Telecommunications Research Institute(ETRI) grant funded by the Korean government (25ZB1100, Core Technology Research for Self-Improving Integrated Artificial Intelligence System), and Electronics and Telecommunications Research Institute(ETRI) grant funded by the Korean government [N01230878, Development of Beyond X-verse Core Technology for Hyper-realistic interactions by Synchronizing the Real World and Virtual Space].

## Author contributions

S.W.L., J.H.L and J.S. conceived and designed the study. J.S. implemented the behavioral task and ran the fMRI study. J.H.L implemented the initial task controller and the simulation framework. J.S. designed computational models and analyzed the data. S.W.L and J.S. wrote the paper. S.W.L, J.H.L and J.S. reviewed, revised and edited the paper. All authors approved the final version for submission.

## Additional information

**Supplementary information** is available for this paper.

**Correspondence** and requests for materials should be addressed to S.W.L.

## Supplementary Figures

**Extended Data Figure 1, Related to Figure 3.**
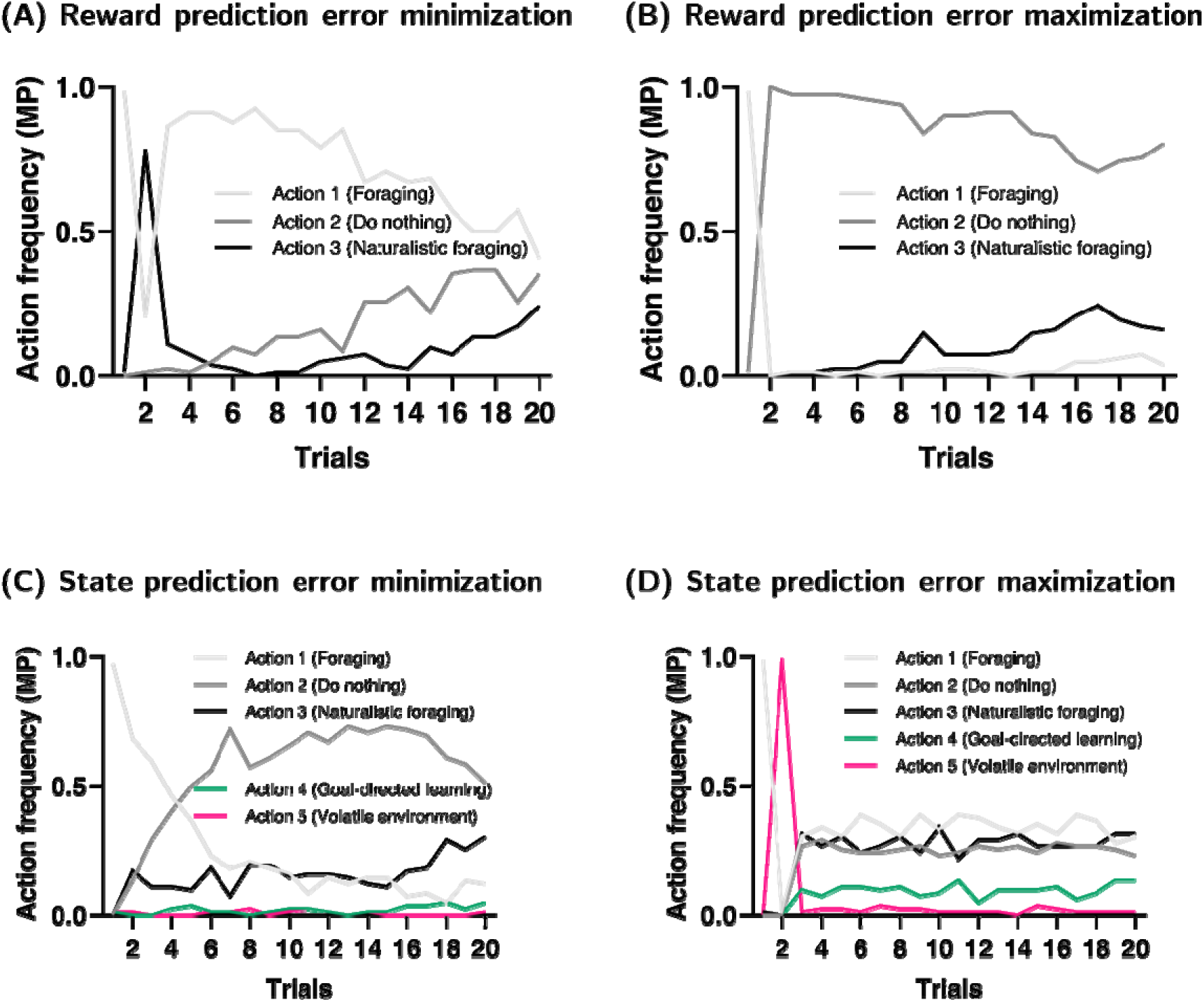
The action sequences of subject-specific meta-prediction models (MPs) after training. The y-axis represented the frequency of actions across 82 MPs. In trial 1, all MPs were set to the ‘Foraging’ (action 1) action. (A) MPs minimizing reward prediction error selected the ‘Naturalistic foraging’ action at trial 2, followed by consistent ‘Foraging’ (action 1) actions, reducing reward discrepancies between optimal and suboptimal choices. (B) MPs maximizing reward prediction error favored the ‘Do nothing’ (action 2) actions from trial two onward, maintaining a steady reward discrepancy between optimal and suboptimal choices. (C) MPs minimizing state prediction error maintained the initial low volatility (state transition probability = [0.9, 0.1]) by ignoring the ‘Volatile environment’ (action 5) action and frequently chose the ‘Goal-directed learning’ (action 4) action to reinforce goal-directed behaviors in the HP. (D) MPs maximizing state prediction error quickly shifted environmental volatility to a high stochastic level (state transition probability = [0.5, 0.5]) and kept it by using the ‘Volatile environment’ (action 5) action only in trial 2. Moreover, MPs used the ‘Goal-directed learning’ (action 4) action less than other actions, promoting a shift in HPs from goal-directed to more habitual behaviors.

**Extended Data Figure 2, Related to Figure 3.**
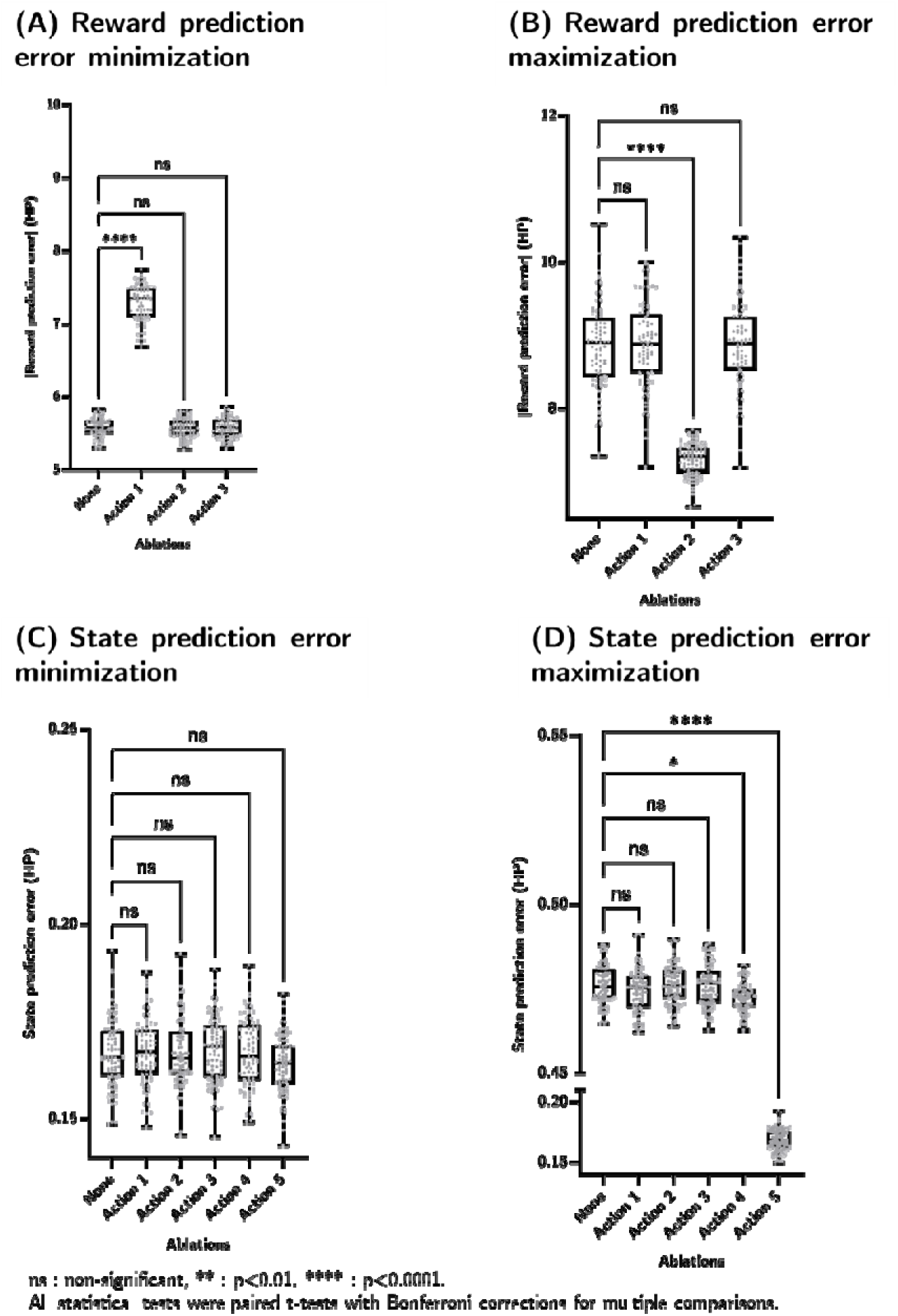
The effect of action ablation in meta-prediction models (MPs) on HP behavior. On the x-axis, ‘None’ represented fully intact MPs, while ‘Action #’ denoted MPs with specific action removed. (A) MPs minimizing reward prediction error showed significantly impaired performance in reward error control when the key action ‘Foraging’ (action 1) was ablated. (B) Removing the ‘Do nothing’ action (action 2) in the MPs maximizing reward prediction error led to a significant performance reduction. (C) The action ablation effects on the MPs minimizing state prediction error were non-significant. (D) For state prediction error maximization, ablation of the ‘Volatile environment’ action (action 5) significantly impaired performance. Ablating ‘Goal-directed learning’ (action 4) also reduced state prediction error control. Asterisks (*) indicate significance levels: * p<0.05, ** p<0.01, *** p<0.001, **** p<0.0001 (two-tailed paired t-test with n = 82).

**Extended Data Figure 3, Related to Figure 4.**
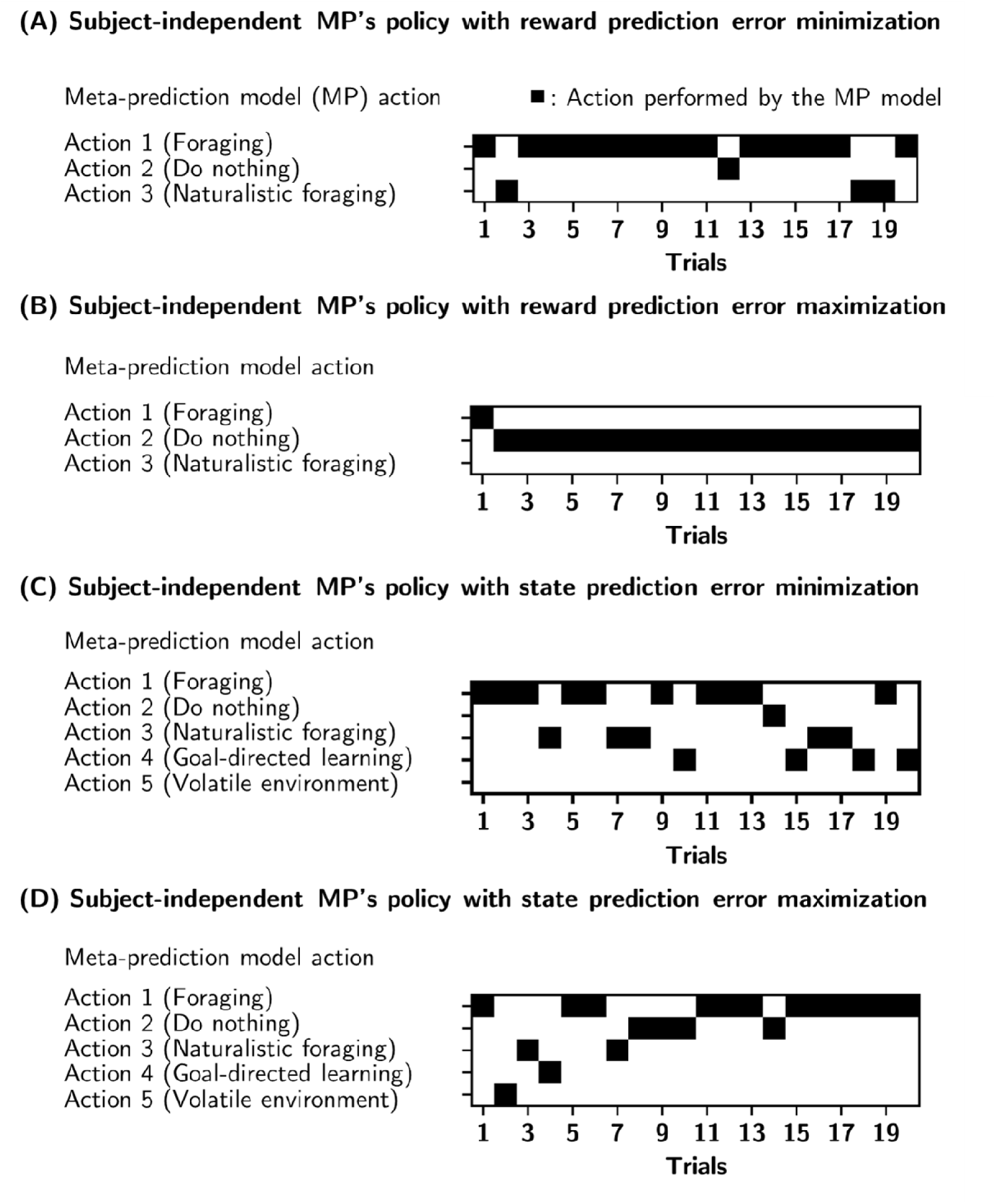
Subject-independent meta-prediction (MP) policies under four conditions: (A) MinR, (B) MaxR, (C) MinS and (D) MaxS. Black boxes mark the actions taken by the MP model.

**Extended Data Figure 4, Related to Figure 4.**
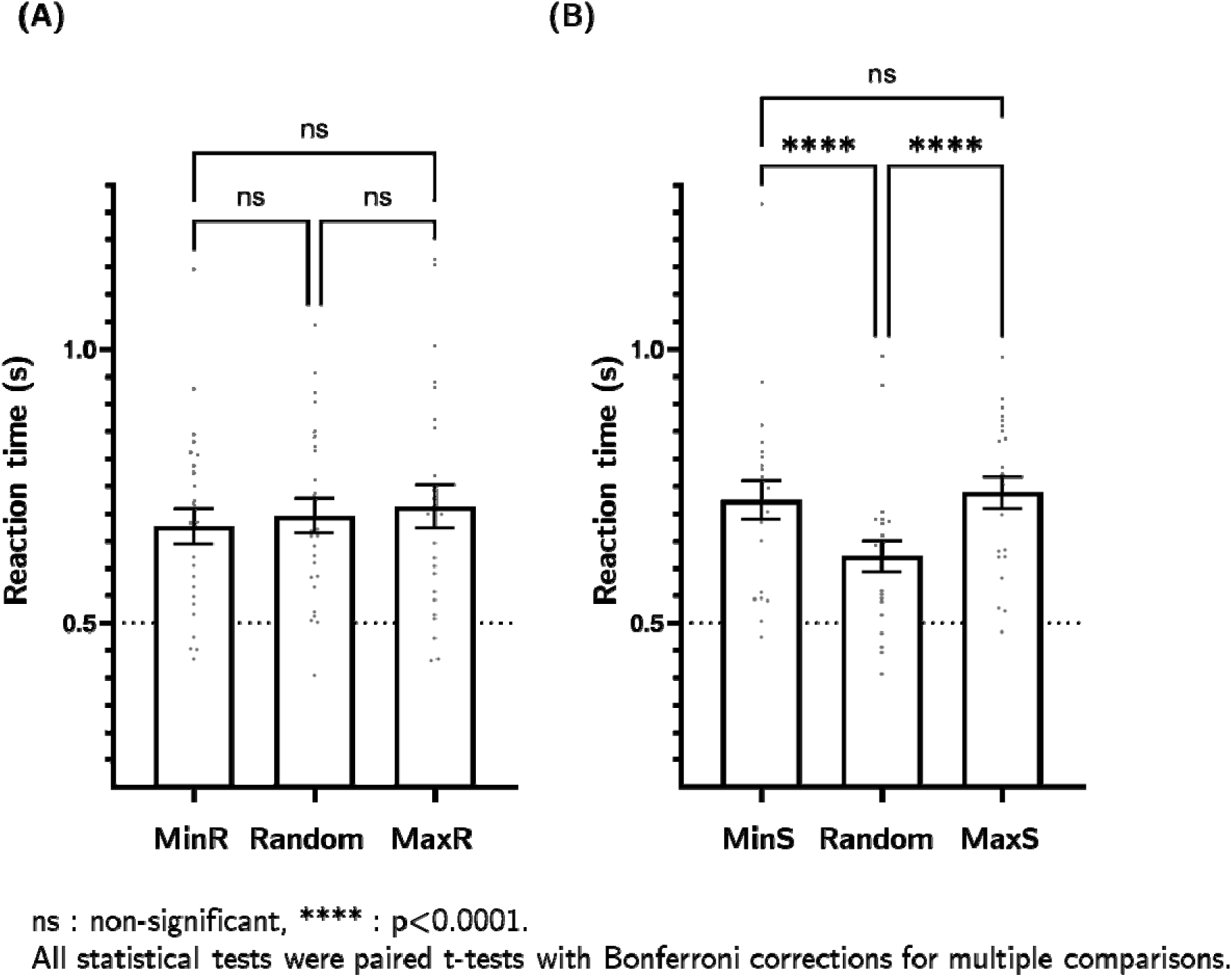
The meta-prediction models (MPs)’ effect on the reaction time of human subjects. (i) Subject-wise (n=26) averaged reaction time under MP conditions for controlling reward prediction errors. The x-axis labels represent: ‘MinR’ = reward prediction error minimization, ‘Random’ = pseudo-random actions, and ‘MaxR’ = reward prediction error maximization. (ii) Subject-wise (n=23) averaged reaction time under MP conditions for controlling state prediction errors. The x-axis labels represent: ‘MinS’ = state prediction error minimization, ‘Random’ = pseudo-random actions, and ‘MaxS’ = state prediction error maximization. The error bars represent SEM, while dots represent the individual subject’s data (n=26 for (i), n=23 for (ii)). Asterisks (*) indicate significance levels: *** p<0.001 (two-tailed paired t-test with n = 26 for (i) and n = 23 for (ii)).

**Extended Data Figure 5, Related to Figure 5.**
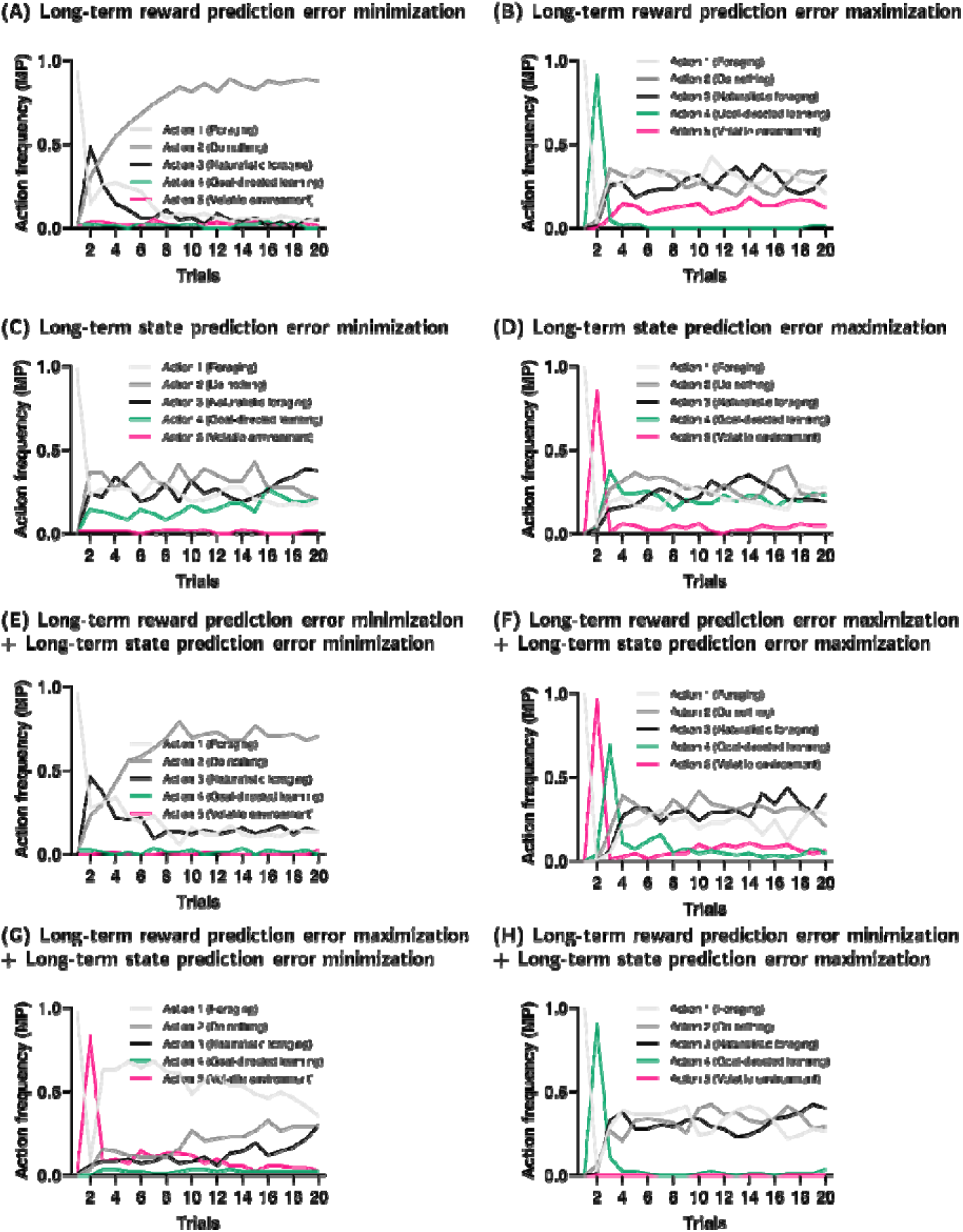
The action sequences of subject-specific meta-prediction models (MP) are trained for controlling long-term prediction errors. The y-axis represented the frequency of actions across 82 MPs. In trial 1, all MPs were set to the ‘Foraging’ (action 1) action. (A-D) MPs controlling a single type of prediction error displayed distinct action profiles: (A) a ‘Do nothing’ (action 2) plateau; (B) a ‘Goal-directed learning’ (action 5) peak at trial 2; (C) ‘Volatile environment’ (action 5) avoidance; and (D) a ‘Volatile environment’ (action 5) peak at trial 2. (E-H) MPs controlling both prediction errors exhibited combined action profiles from relevant single-error MPs. (E) showed a mix of (B) ‘‘Goal-directed learning’ (action 4) peaked at trial 2 and (D) ‘Volatile environment’ (action 5) peaked at trial 3. (F) combined (A) ‘Do nothing’ (action 2) plateau and (C) ‘Volatile environment’ (action 5) avoidance. (G) combined partial (B) ‘Do nothing’ (action 2) plateau with (D) ‘Volatile environment’ (action 5) peak at trial 2. (H) combined (B) ‘Goal-directed learning’ (action 4) peak at trial 2 and (C) ‘Volatile environment’ (action 5) avoidance

**Extended Data Figure 6, Related to Figure 5.**
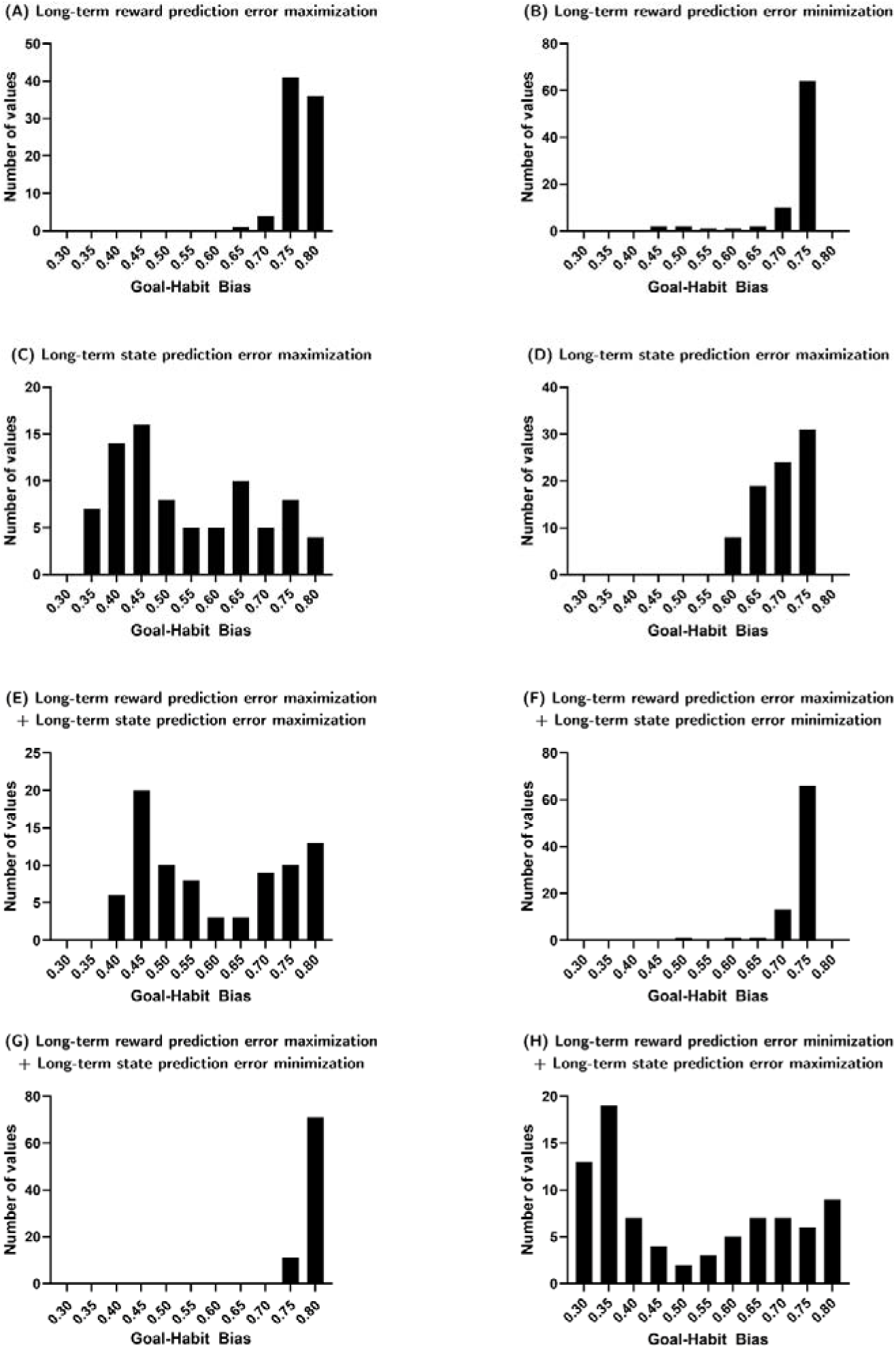
The goal-habit bias of Human Prediction models (HPs) with trained subject-specific Meta Prediction models (MPs) for long-term prediction error controls. (A-D) The goal-habit bias of HPs with MPs controlling only one type of prediction error. Only the MPs maximizing long-term state prediction error (C) stretched the goal-habit bias of HPs toward habitual direction. (E-H) The goal-habit bias of HPs with MPs controlling both types of prediction errors. The MPs that maximize the long-term prediction errors (E, H) showed the goal-habit bias separation among HPs.

**Extended Data Figure 7, Related to Figure 5.**
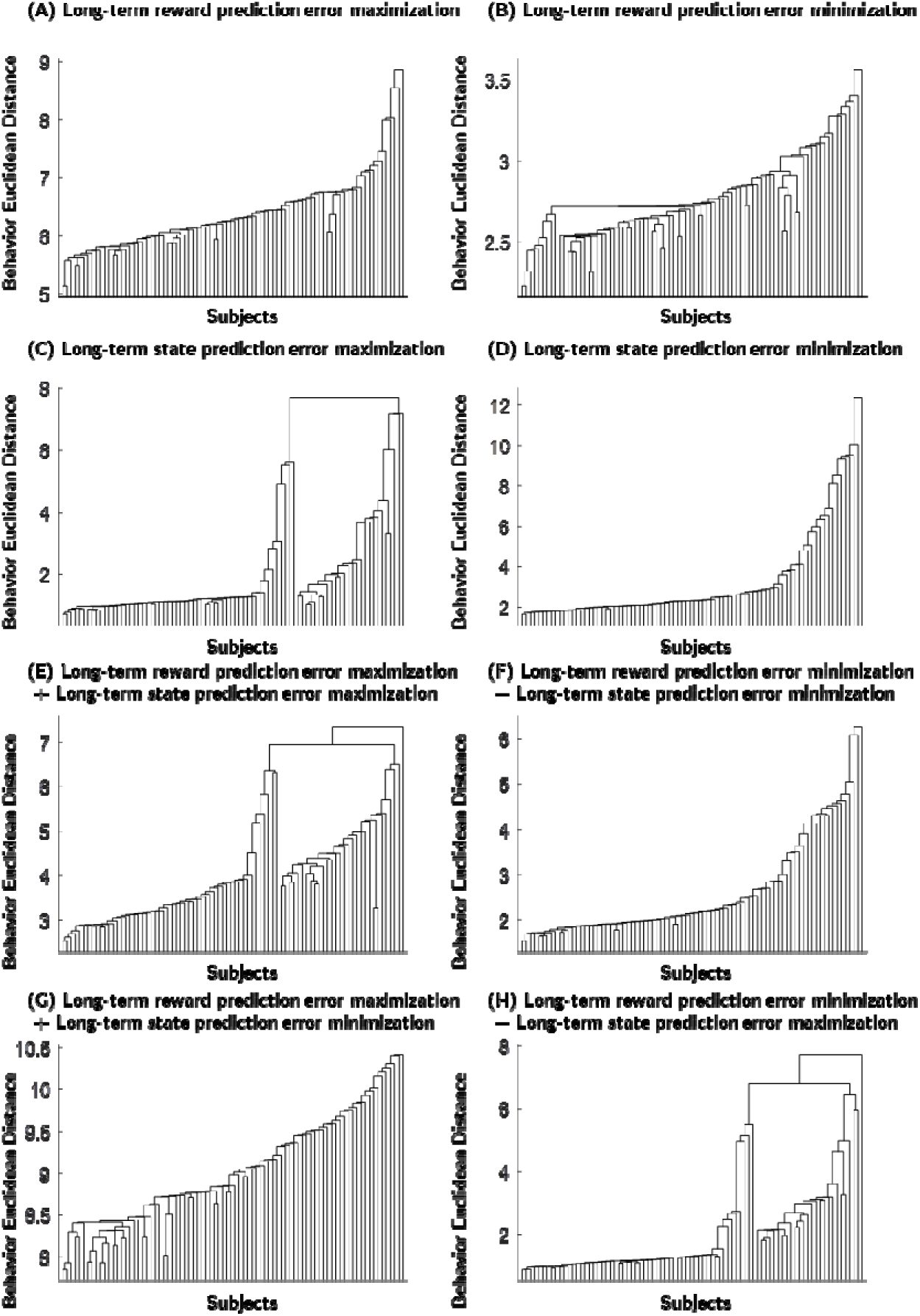
The distance between the Human Prediction model (HP) ‘s simulated behavior from the whole set of trained subject-specific Meta Prediction models(MPs). The x-axis was automatically sorted for visualization. The HP behaviors under MPs with the long-term state prediction error maximizations (C, E, H) showed two clusters in the dendrogram, while other MPs (A, B, D, F, G) did not show the separations.

**Extended Data Figure 8, Related to Figure 5.**
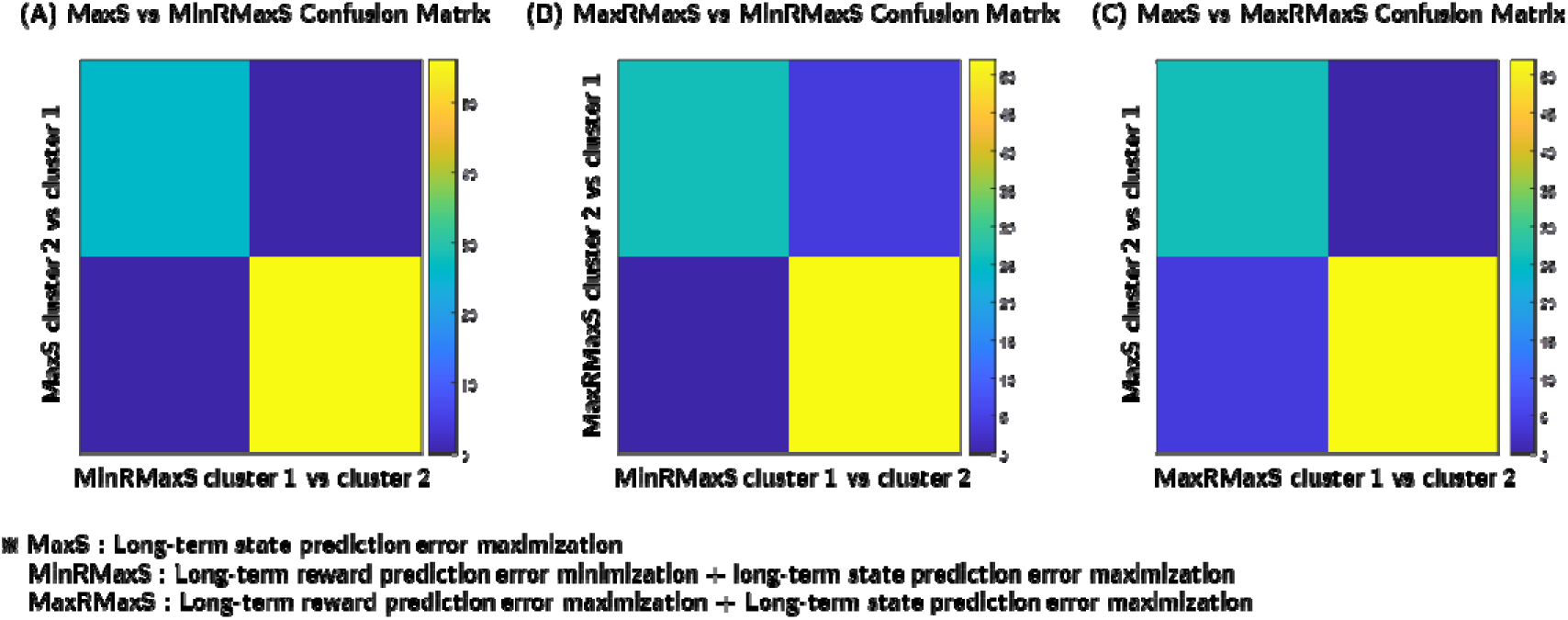
The confusion matrices between Human Prediction model (HP) clusterings under 3 Meta Prediction models (MPs; The long-term state prediction error maximization only; MaxS, The long-term reward prediction error maximization + the long-term state prediction minimization; MaxRMaxS, The long-term reward prediction error minimization + the long-term state prediction minimization; MinRMaxS). The HP clusters were replicated in independent classifications with each MP.

**Extended Data Figure 9, Related to Figure 5.**
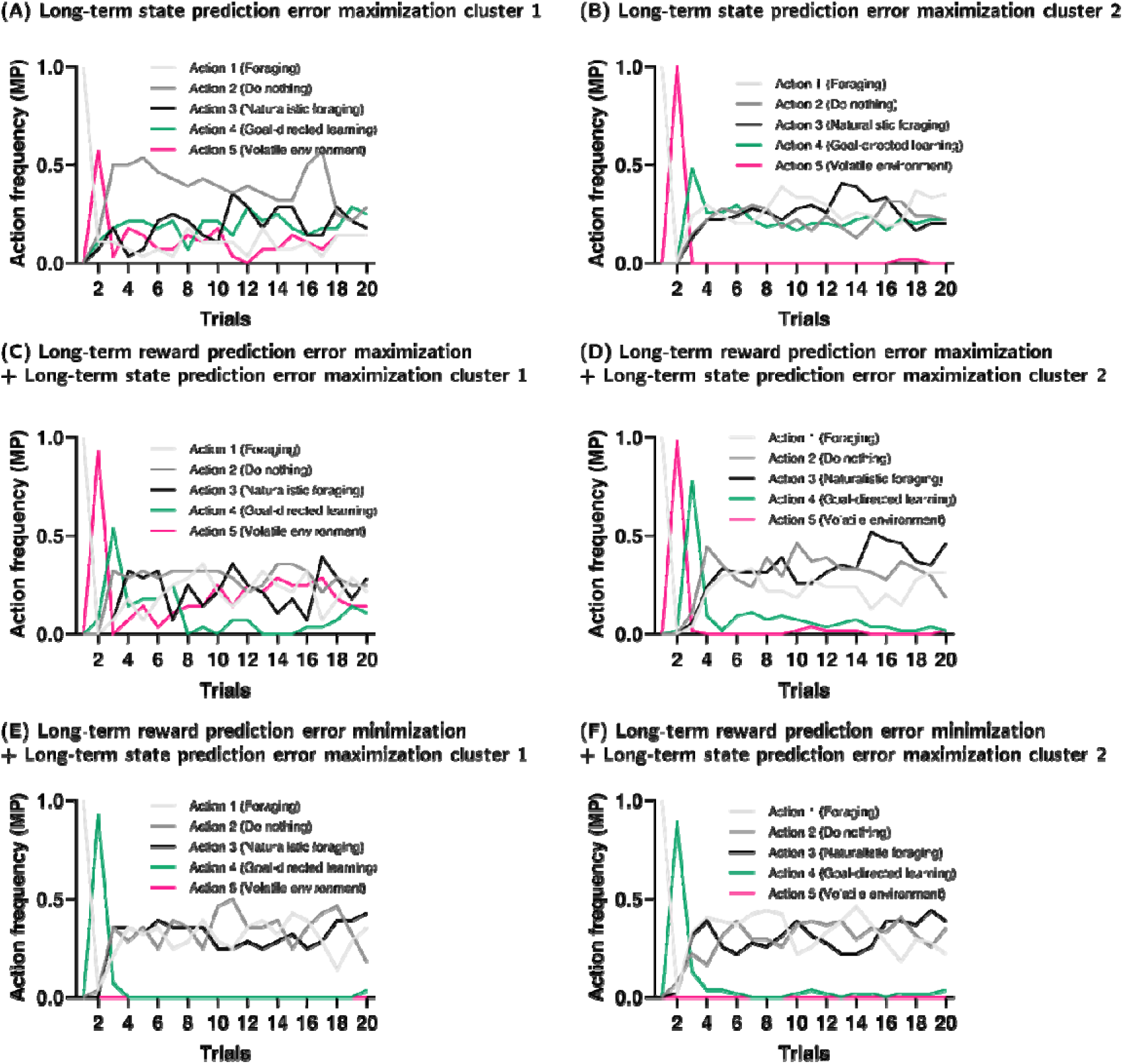
Optimized action sequences of subject-specific meta-prediction models (MPs) controlling long-term prediction errors within each cluster. The y-axis represented action frequency across 82 MPs in a specific trial. All MPs were set to ‘Foraging’ (action 1) in trial 1. (A, B) MPs maximizing only the long-term state prediction error. Both clusters exhibited a prominent ‘Volatile environment’ (action 5) peak in trial 2. From trial three onward, clusters diverged: cluster 1 MPs (A) favored ‘Do nothing’ (action 2) over other actions in most of the remaining trials, while cluster 2 MPs (B) mostly selected ‘Goal-directed learning’ (action 4) in trial 3, then made random actions, excluding ‘Volatile environment’ (action 5), after trial 3. (C, D) MPs maximizing both long-term reward prediction error and long-term state prediction error. The MPs followed a mixed policy, combining the behavior of long-term state prediction error MPs in the same cluster with the policy of long-term reward prediction error maximization MPs (see Extended Data Fig. 5B). (E, F) MPs minimizing long-term reward prediction error and maximizing long-term state prediction error. These MPs followed a mixed policy, combining the behavior of long-term state prediction error MPs in the same cluster with that of long-term reward prediction error minimization MPs (see Extended Data Fig. 5A).

## Supplementary Data

**Table S1.** Parameter values of human prediction models in Meta-prediction training, Related to Figure 1C.

| parameter<br>subject | 1 | 2 | 3 | 4 | 5 | 6 |
| --- | --- | --- | --- | --- | --- | --- |
| 1 | 0.614434 | 0.077846 | 0.485981 | 6.358767 | 0.02833 | 0.199619 |
| 2 | 0.768861 | 0.303617 | 0.121948 | 9.841435 | 0.163633 | 0.079116 |
| 3 | 0.789506 | 0.348951 | 0.104459 | 7.674034 | 0.151593 | 0.069762 |
| 4 | 0.409728 | 0.106105 | 0.115738 | 3.459498 | 0.109123 | 0.100876 |
| 5 | 0.526636 | 0.289645 | 0.100623 | 9.439895 | 0.092528 | 0.141934 |
| 6 | 0.606038 | 0.342951 | 0.111231 | 9.076079 | 0.190424 | 0.131374 |
| 7 | 0.760805 | 0.257565 | 0.111405 | 8.585745 | 0.24738 | 0.049113 |
| 8 | 0.547765 | 0.097489 | 0.10841 | 3.83039 | 0.267223 | 0.023925 |
| 9 | 0.655238 | 0.348549 | 0.141602 | 8.84792 | 0.119685 | 0.097181 |
| 10 | 0.662204 | 0.035764 | 0.113472 | 9.770955 | 0.232971 | 0.176474 |
| 11 | 0.44467 | 0.047784 | 0.106051 | 0.493613 | 0.131951 | 0.198862 |
| 12 | 0.310721 | 0.051232 | 0.329584 | 0.133219 | 0.070455 | 0.191334 |
| 13 | 0.571157 | 0.322415 | 0.12557 | 9.499509 | 0.042897 | 0.07735 |
| 14 | 0.43934 | 0.095588 | 0.116781 | 0.145777 | 0.113934 | 0.187598 |
| 15 | 0.31271 | 0.010084 | 0.128219 | 6.354347 | 0.075998 | 0.199572 |
| 16 | 0.310466 | 0.015333 | 0.100762 | 0.100675 | 0.034124 | 0.178554 |
| 17 | 0.388709 | 0.023578 | 0.110374 | 0.874238 | 0.092507 | 0.090436 |
| 18 | 0.431981 | 0.01841 | 9.12556 | 5.438772 | 0.073431 | 0.198329 |
| 19 | 0.407993 | 0.048459 | 0.117829 | 0.176122 | 0.200654 | 0.086718 |
| 20 | 0.779709 | 0.341282 | 0.130899 | 9.993625 | 0.133518 | 0.086143 |
| 21 | 0.752672 | 0.346166 | 0.132993 | 7.2859 | 0.107888 | 0.01332 |
| 22 | 0.322062 | 0.024744 | 0.106951 | 0.124803 | 0.165586 | 0.068983 |
| 23 | 0.446239 | 0.246659 | 0.779179 | 6.856203 | 0.121264 | 0.091161 |
| 24 | 0.371982 | 0.228268 | 2.823635 | 5.559278 | 0.065565 | 0.199967 |
| 25 | 0.381629 | 0.111274 | 0.109168 | 6.159796 | 0.049914 | 0.128051 |
| 26 | 0.788474 | 0.340892 | 0.110013 | 8.002945 | 0.052318 | 0.100677 |
| 27 | 0.321857 | 0.017022 | 0.111729 | 1.000458 | 0.178763 | 0.014052 |
| 28 | 0.300924 | 0.029971 | 9.905311 | 6.714189 | 0.061418 | 0.034508 |
| 29 | 0.324384 | 0.09861 | 9.9952 | 0.115685 | 0.126445 | 0.199985 |
| 30 | 0.537777 | 0.296909 | 0.11631 | 2.262599 | 0.088132 | 0.195528 |
| 31 | 0.592012 | 0.338338 | 0.110825 | 9.999963 | 0.148137 | 0.177133 |
| 32 | 0.301583 | 0.342705 | 9.146814 | 9.922909 | 0.010052 | 0.010093 |
| 33 | 0.755449 | 0.314772 | 0.170361 | 8.759698 | 0.081446 | 0.033905 |
| 34 | 0.776697 | 0.346732 | 9.997266 | 9.987646 | 0.059832 | 0.199926 |
| 35 | 0.605991 | 0.186202 | 0.432683 | 6.133691 | 0.010068 | 0.010406 |
| 36 | 0.302178 | 0.01087 | 0.134902 | 1.309548 | 0.045499 | 0.199933 |
| 37 | 0.531646 | 0.095563 | 0.111375 | 3.536236 | 0.092553 | 0.021334 |
| 38 | 0.637308 | 0.342439 | 0.109019 | 7.637068 | 0.221664 | 0.092096 |
| 39 | 0.59083 | 0.014004 | 0.111107 | 6.30222 | 0.172549 | 0.141236 |
| 40 | 0.362292 | 0.056831 | 1.003234 | 0.194567 | 0.065983 | 0.01008 |
| 41 | 0.300063 | 0.010425 | 0.109499 | 0.735191 | 0.249828 | 0.166104 |
| 42 | 0.39453 | 0.027757 | 0.290048 | 1.50953 | 0.057533 | 0.077469 |
| 43 | 0.415871 | 0.06227 | 0.13131 | 2.027751 | 0.046044 | 0.146878 |
| 44 | 0.786126 | 0.095856 | 0.109239 | 9.074241 | 0.15545 | 0.117417 |
| 45 | 0.303426 | 0.331936 | 0.513855 | 7.642088 | 0.046049 | 0.19903 |
| 46 | 0.712641 | 0.344217 | 4.342648 | 9.410404 | 0.135017 | 0.199966 |
| 47 | 0.302296 | 0.115472 | 9.737553 | 2.706059 | 0.09124 | 0.15917 |
| 48 | 0.403089 | 0.032476 | 0.110356 | 4.048887 | 0.021734 | 0.083046 |
| 49 | 0.788937 | 0.082713 | 0.166061 | 8.764681 | 0.094145 | 0.199098 |
| 50 | 0.635781 | 0.337506 | 0.113893 | 9.691597 | 0.184215 | 0.095825 |
| 51 | 0.627248 | 0.31874 | 0.112056 | 8.719899 | 0.115659 | 0.12472 |
| 52 | 0.690221 | 0.130822 | 0.111413 | 9.197222 | 0.137439 | 0.057715 |
| 53 | 0.761633 | 0.033442 | 0.273668 | 9.813905 | 0.032257 | 0.010046 |
| 54 | 0.315846 | 0.057795 | 0.112027 | 0.112444 | 0.370857 | 0.103709 |
| 55 | 0.329614 | 0.264606 | 0.183554 | 2.144173 | 0.062925 | 0.17437 |
| 56 | 0.485611 | 0.013314 | 0.15194 | 0.101912 | 0.30808 | 0.090112 |
| 57 | 0.35893 | 0.329849 | 0.104179 | 9.616879 | 0.10953 | 0.112101 |
| 58 | 0.414693 | 0.33331 | 0.165218 | 9.657266 | 0.085795 | 0.199955 |
| 59 | 0.50646 | 0.118649 | 0.116518 | 2.924663 | 0.082591 | 0.179123 |
| 60 | 0.705979 | 0.33874 | 0.103961 | 9.991226 | 0.095651 | 0.126693 |
| 61 | 0.475085 | 0.108457 | 0.116342 | 3.730337 | 0.134566 | 0.113445 |
| 62 | 0.430969 | 0.347205 | 0.113076 | 9.993661 | 0.301218 | 0.13597 |
| 63 | 0.364989 | 0.335778 | 0.106356 | 9.525257 | 0.325967 | 0.160961 |
| 64 | 0.313824 | 0.066376 | 0.108005 | 0.746548 | 0.337433 | 0.152106 |
| 65 | 0.305608 | 0.023424 | 0.110639 | 0.100247 | 0.16997 | 0.095547 |
| 66 | 0.478126 | 0.015675 | 0.113917 | 0.104709 | 0.370248 | 0.012558 |
| 67 | 0.312882 | 0.265409 | 0.110839 | 6.826111 | 0.355819 | 0.022437 |
| 68 | 0.408621 | 0.34813 | 0.116268 | 9.545918 | 0.132775 | 0.102069 |
| 69 | 0.737792 | 0.349941 | 0.110716 | 9.088162 | 0.329604 | 0.109003 |
| 70 | 0.300125 | 0.328244 | 0.107688 | 9.979268 | 0.400769 | 0.077915 |
| 71 | 0.324178 | 0.020972 | 0.116942 | 1.075751 | 0.118785 | 0.032027 |
| 72 | 0.328015 | 0.012474 | 0.107297 | 2.428303 | 0.150773 | 0.199184 |
| 73 | 0.689799 | 0.144685 | 1.92092 | 2.926623 | 0.168029 | 0.159962 |
| 74 | 0.310902 | 0.056944 | 0.376634 | 3.406745 | 0.032164 | 0.067326 |
| 75 | 0.303094 | 0.058424 | 0.119711 | 0.482937 | 0.252542 | 0.010001 |
| 76 | 0.318385 | 0.0385 | 0.107431 | 9.728752 | 0.162256 | 0.110868 |
| 77 | 0.734543 | 0.349841 | 0.115792 | 9.647542 | 0.227691 | 0.140819 |
| 78 | 0.331594 | 0.014504 | 0.1101 | 0.10008 | 0.338421 | 0.191684 |
| 79 | 0.596621 | 0.32534 | 9.936829 | 9.266734 | 0.220292 | 0.010664 |
| 80 | 0.777138 | 0.349217 | 9.979405 | 8.676161 | 0.031022 | 0.197258 |
| 81 | 0.54892 | 0.308085 | 0.119453 | 7.895054 | 0.117192 | 0.197752 |
| 82 | 0.369014 | 0.013708 | 0.110438 | 3.257821 | 0.063261 | 0.141257 |

Parameter: 1- the threshold for defining zero state prediction error (SPE tolerance), 2- learning rate for the estimate of absolute reward prediction error, 3- rate of the habit-to-goal-directed transition, 4-rate of the goal-directed-to-habit transition, 5- inverse softmax temperature, and 6- learning rate of the goal-directed and the habitual learning systems, respectively.

**Table S2.**
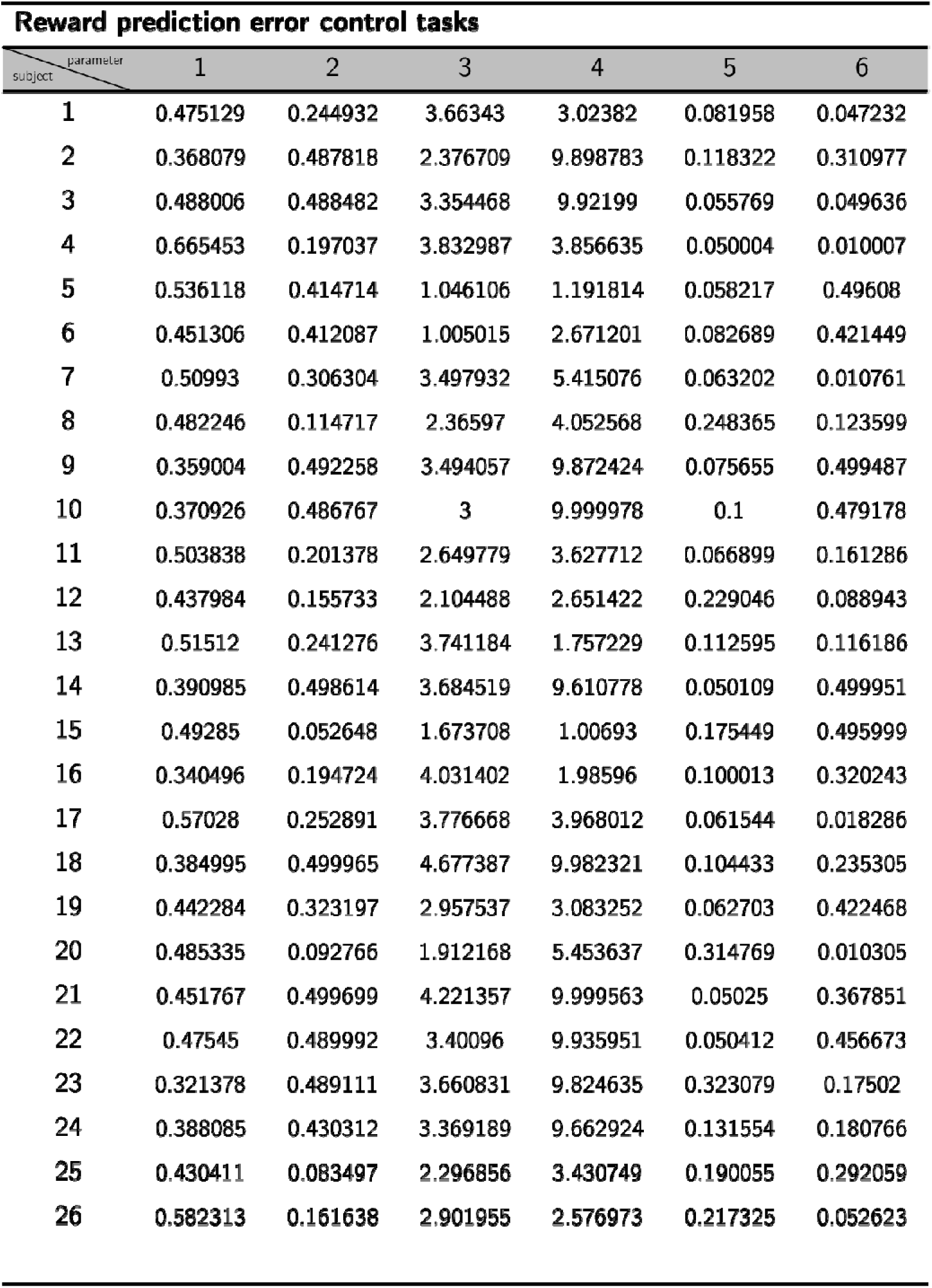
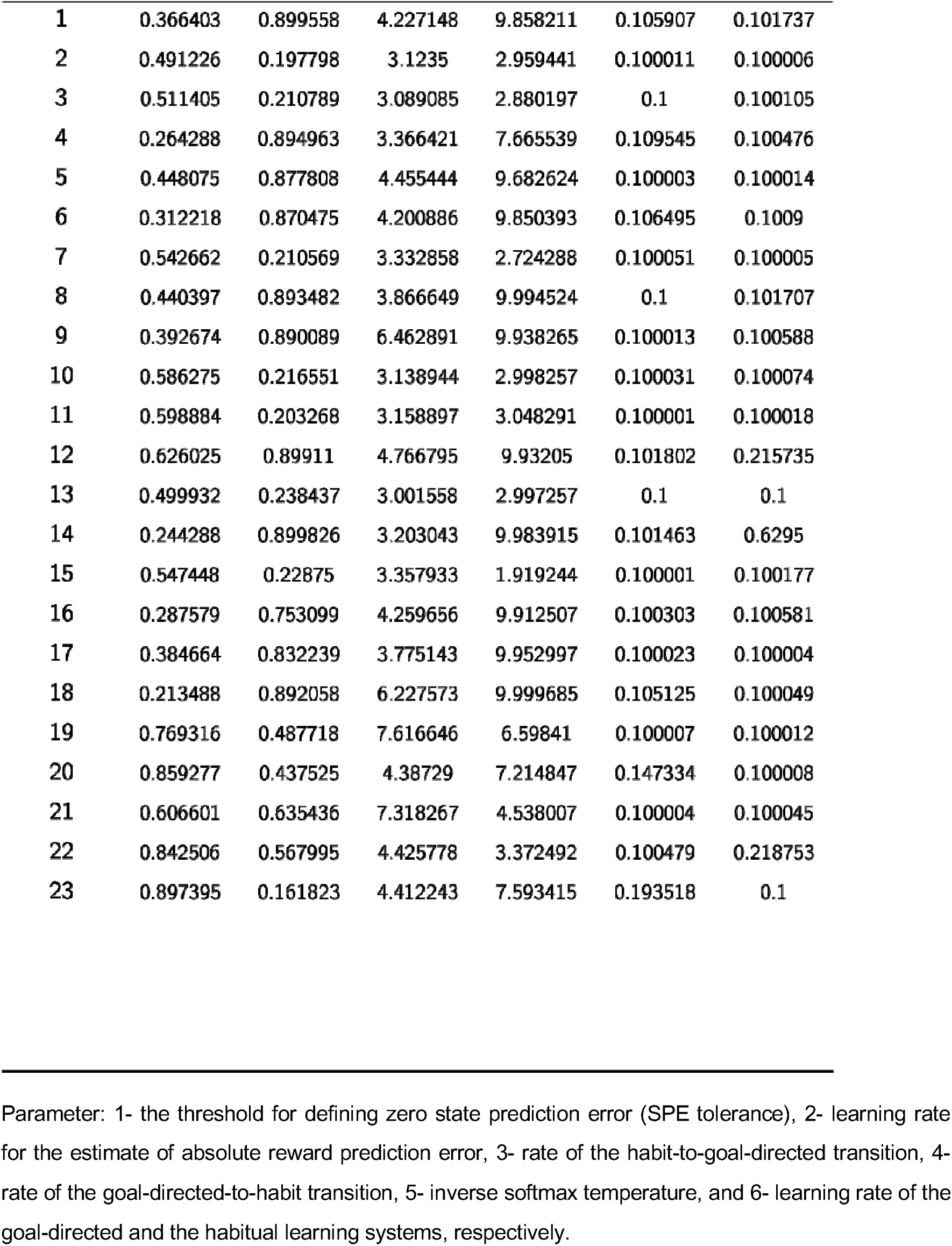
Estimated parameter values of the prefrontal arbitration model, Related to Figure 4.

| Reward prediction error control tasks |  |  |  |  |  |  |
| --- | --- | --- | --- | --- | --- | --- |
| subject \ parameter | 1 | 2 | 3 | 4 | 5 | 6 |
| 1 | 0.475129 | 0.244932 | 3.66343 | 3.02382 | 0.081958 | 0.047232 |
| 2 | 0.368079 | 0.487818 | 2.376709 | 9.898783 | 0.118322 | 0.310977 |
| 3 | 0.488006 | 0.488482 | 3.354468 | 9.92199 | 0.055769 | 0.049636 |
| 4 | 0.665453 | 0.197037 | 3.832987 | 3.856635 | 0.050004 | 0.010007 |
| 5 | 0.536118 | 0.414714 | 1.046106 | 1.191814 | 0.058217 | 0.49608 |
| 6 | 0.451306 | 0.412087 | 1.005015 | 2.671201 | 0.082689 | 0.421449 |
| 7 | 0.50993 | 0.306304 | 3.497932 | 5.415076 | 0.063202 | 0.010761 |
| 8 | 0.482246 | 0.114717 | 2.36597 | 4.052568 | 0.248365 | 0.123599 |
| 9 | 0.359004 | 0.492258 | 3.494057 | 9.872424 | 0.075655 | 0.499487 |
| 10 | 0.370926 | 0.486767 | 3 | 9.999978 | 0.1 | 0.479178 |
| 11 | 0.503838 | 0.201378 | 2.649779 | 3.627712 | 0.066899 | 0.161286 |
| 12 | 0.437984 | 0.155733 | 2.104488 | 2.651422 | 0.229046 | 0.088943 |
| 13 | 0.51512 | 0.241276 | 3.741184 | 1.757229 | 0.112595 | 0.116186 |
| 14 | 0.390985 | 0.498614 | 3.684519 | 9.610778 | 0.050109 | 0.499951 |
| 15 | 0.49285 | 0.052648 | 1.673708 | 1.00693 | 0.175449 | 0.495999 |
| 16 | 0.340496 | 0.194724 | 4.031402 | 1.98596 | 0.100013 | 0.320243 |
| 17 | 0.57028 | 0.252891 | 3.776668 | 3.968012 | 0.061544 | 0.018286 |
| 18 | 0.384995 | 0.499965 | 4.677387 | 9.982321 | 0.104433 | 0.235305 |
| 19 | 0.442284 | 0.323197 | 2.957537 | 3.083252 | 0.062703 | 0.422468 |
| 20 | 0.485335 | 0.092766 | 1.912168 | 5.453637 | 0.314769 | 0.010305 |
| 21 | 0.451767 | 0.499699 | 4.221357 | 9.999563 | 0.05025 | 0.367851 |
| 22 | 0.47545 | 0.489992 | 3.40096 | 9.935951 | 0.050412 | 0.456673 |
| 23 | 0.321378 | 0.489111 | 3.660831 | 9.824635 | 0.323079 | 0.17502 |
| 24 | 0.388085 | 0.430312 | 3.369189 | 9.662924 | 0.131554 | 0.180766 |
| 25 | 0.430411 | 0.083497 | 2.296856 | 3.430749 | 0.190055 | 0.292059 |
| 26 | 0.582313 | 0.161638 | 2.901955 | 2.576973 | 0.217325 | 0.052623 |

| State prediction error control tasks |  |  |  |  |  |  |
| --- | --- | --- | --- | --- | --- | --- |
| subject \ parameter | 1 | 2 | 3 | 4 | 5 | 6 |
| 1 | 0.366403 | 0.899558 | 4.227148 | 9.858211 | 0.105907 | 0.101737 |
| 2 | 0.491226 | 0.197798 | 3.1235 | 2.959441 | 0.100011 | 0.100006 |
| 3 | 0.511405 | 0.210789 | 3.089085 | 2.880197 | 0.1 | 0.100105 |
| 4 | 0.264288 | 0.894963 | 3.366421 | 7.665539 | 0.109545 | 0.100476 |
| 5 | 0.448075 | 0.877808 | 4.455444 | 9.682624 | 0.100003 | 0.100014 |
| 6 | 0.312218 | 0.870475 | 4.200886 | 9.850393 | 0.106495 | 0.1009 |
| 7 | 0.542662 | 0.210569 | 3.332858 | 2.724288 | 0.100051 | 0.100005 |
| 8 | 0.440397 | 0.893482 | 3.866649 | 9.994524 | 0.1 | 0.101707 |
| 9 | 0.392674 | 0.890089 | 6.462891 | 9.938265 | 0.100013 | 0.100588 |
| 10 | 0.586275 | 0.216551 | 3.138944 | 2.998257 | 0.100031 | 0.100074 |
| 11 | 0.598884 | 0.203268 | 3.158897 | 3.048291 | 0.100001 | 0.100018 |
| 12 | 0.626025 | 0.89911 | 4.766795 | 9.93205 | 0.101802 | 0.215735 |
| 13 | 0.499932 | 0.238437 | 3.001558 | 2.997257 | 0.1 | 0.1 |
| 14 | 0.244288 | 0.899826 | 3.203043 | 9.983915 | 0.101463 | 0.6295 |
| 15 | 0.547448 | 0.22875 | 3.357933 | 1.919244 | 0.100001 | 0.100177 |
| 16 | 0.287579 | 0.753099 | 4.259656 | 9.912507 | 0.100303 | 0.100581 |
| 17 | 0.384664 | 0.832239 | 3.775143 | 9.952997 | 0.100023 | 0.100004 |
| 18 | 0.213488 | 0.892058 | 6.227573 | 9.999685 | 0.105125 | 0.100049 |
| 19 | 0.769316 | 0.487718 | 7.616646 | 6.59841 | 0.100007 | 0.100012 |
| 20 | 0.859277 | 0.437525 | 4.38729 | 7.214847 | 0.147334 | 0.100008 |
| 21 | 0.606601 | 0.635436 | 7.318267 | 4.538007 | 0.100004 | 0.100045 |
| 22 | 0.842506 | 0.567995 | 4.425778 | 3.372492 | 0.100479 | 0.218753 |
| 23 | 0.897395 | 0.161823 | 4.412243 | 7.593415 | 0.193518 | 0.1 |

Parameter: 1- the threshold for defining zero state prediction error (SPE tolerance), 2- learning rate for the estimate of absolute reward prediction error, 3- rate of the habit-to-goal-directed transition, 4-rate of the goal-directed-to-habit transition, 5- inverse softmax temperature, and 6- learning rate of the goal-directed and the habitual learning systems, respectively.

**Table S3.**
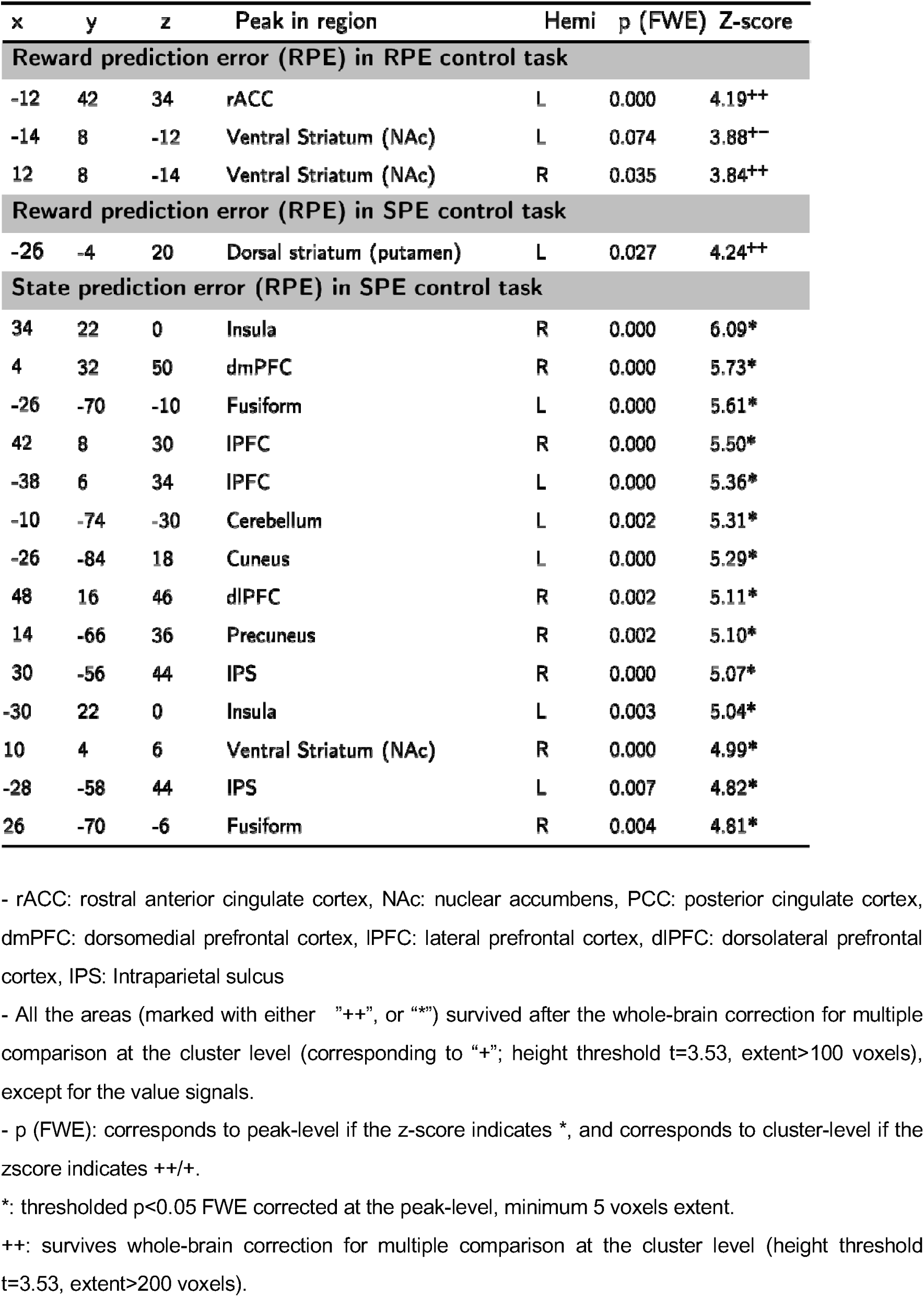
Neural signatures of the habitual and goal-directed systems, Related to Figure 4.

**Table S4.**
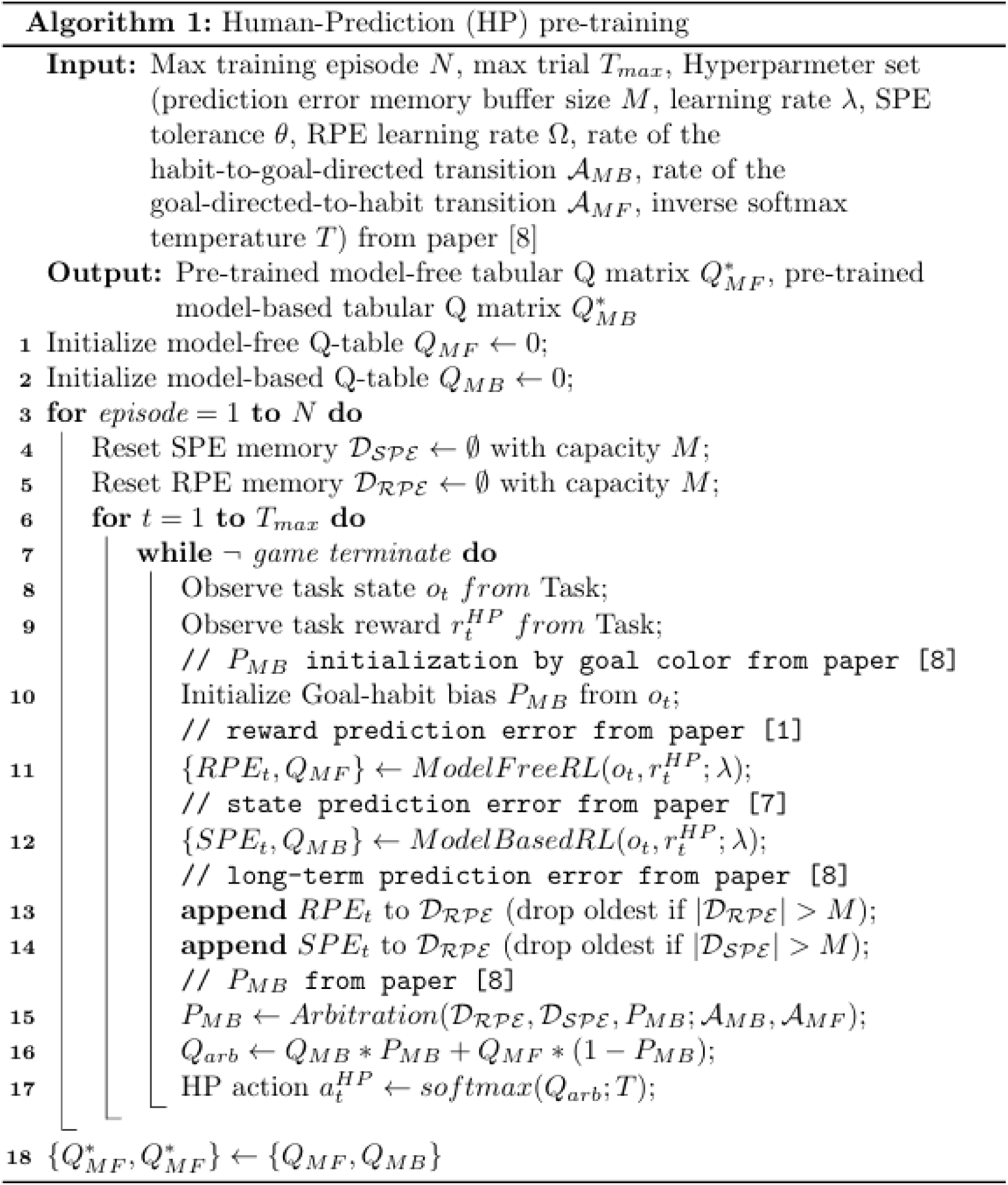
Algorithm for Human-prediction (HP) pre-training, Related to method section.

**Table S5.**
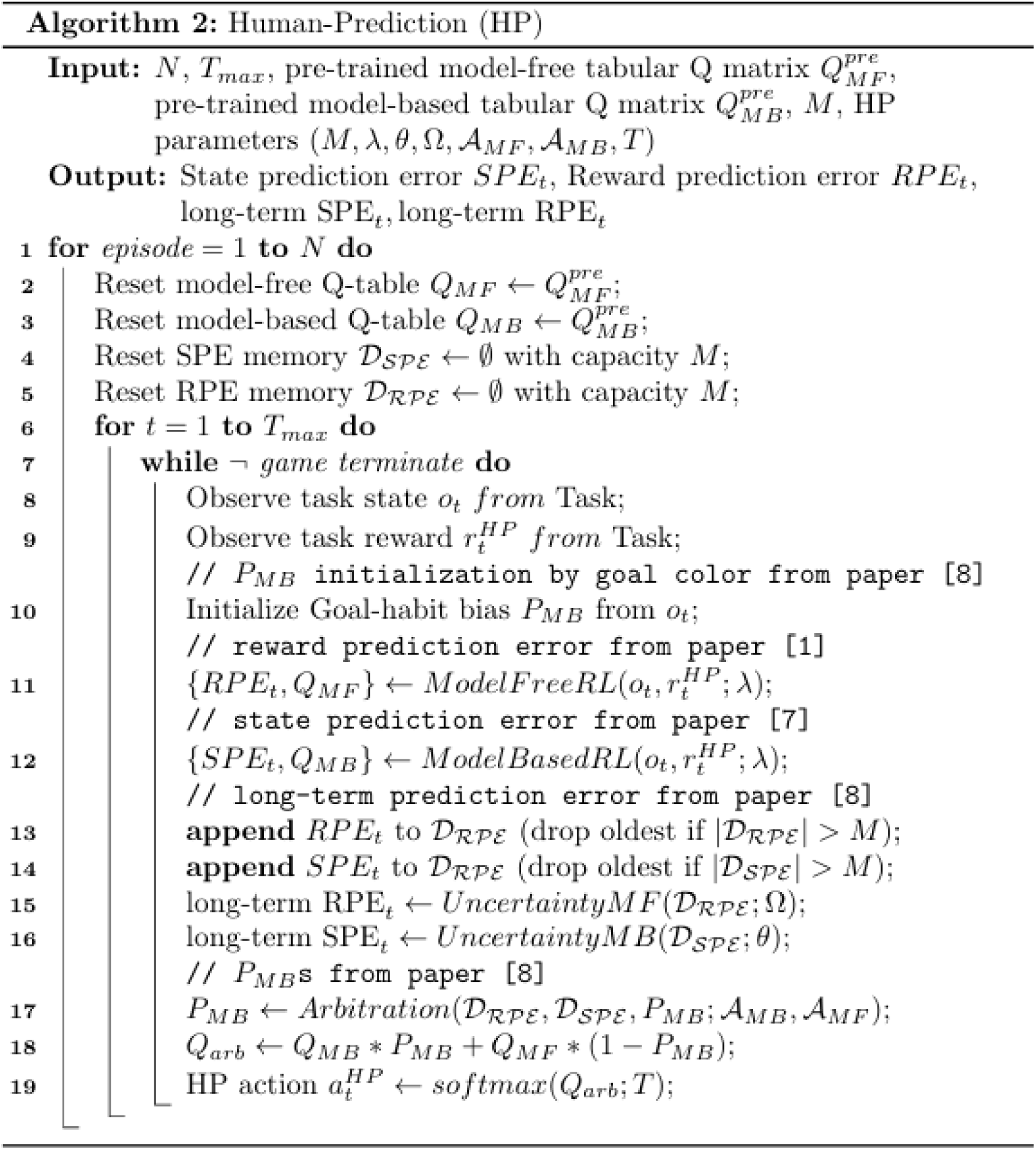
Algorithm for Human-prediction (HP), Related to method section.

**Table S6.**
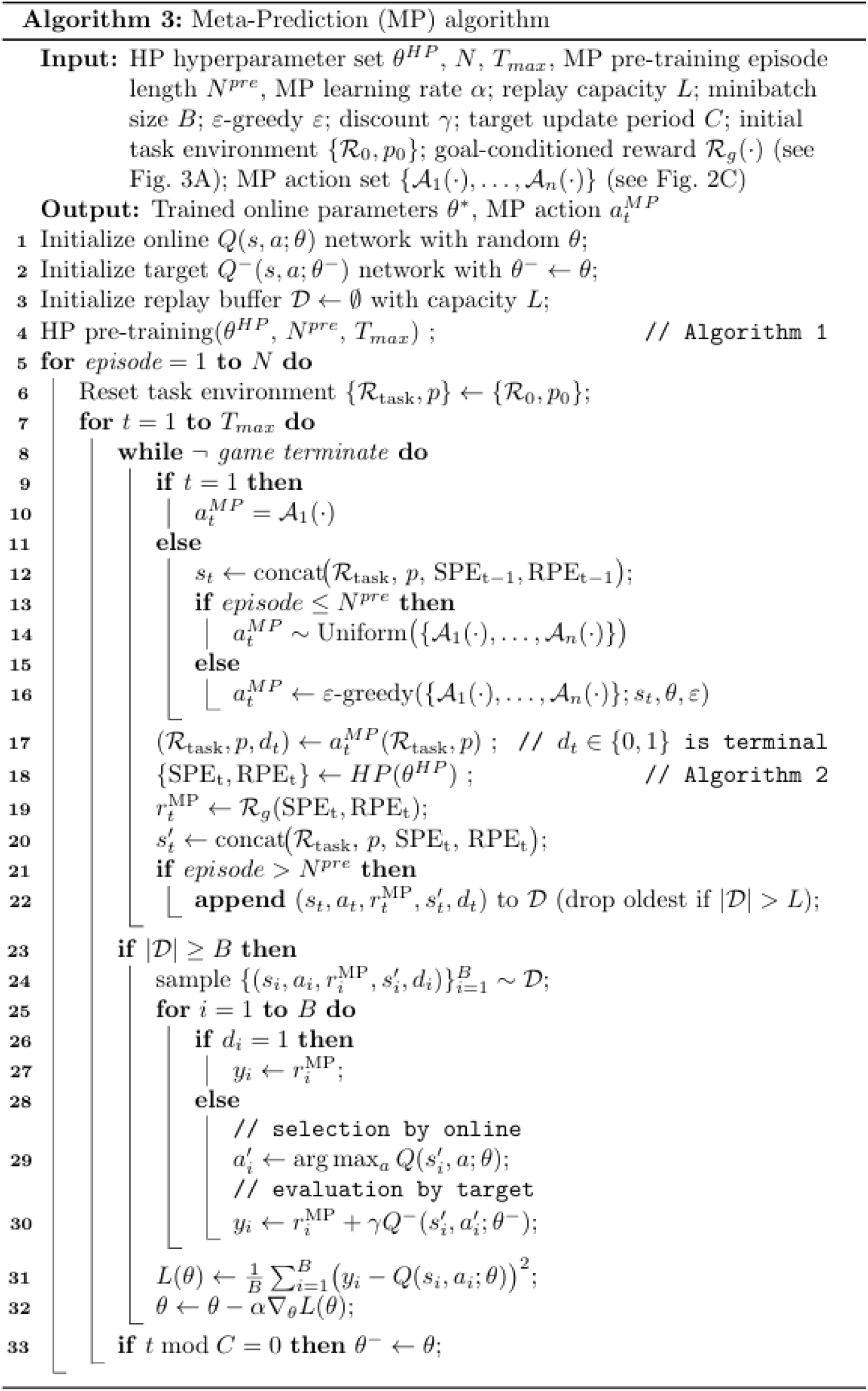
Algorithm for Meta-prediction (MP) pre-training, Related to method section.

